# AnewDDE: An Agentic Drug Discovery Engine for Biomolecular Interaction Modelling and Closed-Loop Design

**DOI:** 10.64898/2026.09.20.752967

**Authors:** Anew Labs Team, Kai Liu

## Abstract

Accurate modelling of biomolecular interactions is fundamental to drug discovery, yet current artificial intelligence (AI) workflows remain fragmented across structure prediction, affinity estimation, molecular design, and experimental decision-making. We introduce AnewDDE, an agentic Drug Discovery Engine that connects these capabilities into a closed-loop system for biomolecular interaction modelling and design. We demonstrate several components of this system: AnewFold delivers superior performance on challenging targets, including antibody–antigen complexes and molecular glues, providing more reliable structural hypotheses together with pocket identification and conformational analysis; AnewDesign achieves a 10.7% success rate in identifying binders with single-digit-nanomolar affinities measured by surface plasmon resonance (SPR) in a representative nanobody discovery campaign, substantially reducing the experimental search burden; AnewAffinity combines accuracy approaching that of free-energy perturbation (FEP) methods with the speed required for high-throughput screening and prioritizes promising transformations; and AnewMind, a large language model (LLM) at the hundred-billion-parameter scale that underwent in-house full-parameter post-training, provides scientific reasoning and absorption, distribution, metabolism, excretion, and toxicity (ADMET) prediction for pharmaceutical research and development (R&D). To systematically evaluate AnewMind, we introduce PharmBench, an internal benchmark designed by pharmaceutical R&D experts to evaluate long-horizon decision-making in drug discovery. Evaluations on ADMET tasks and PharmBench show that AnewMind performs competitively with leading frontier models. Together, these capabilities establish AnewDDE as an integrated engine for addressing challenging therapeutic targets and translating molecular insights into experimentally validated candidates at scale.

## 1 Introduction

Drug discovery relies on the precise prediction and control of biomolecular interactions. Small molecules must recognize target pockets with compatible geometry and energetics, while antibodies must selectively engage functional epitopes. Translating these molecular interactions into viable therapeutics is therefore a complex, multi-dimensional design challenge.

In practice, the objective is rarely a single isolated prediction. Therapeutic design requires an integrated chain of decisions linking structural hypotheses, binding mechanisms, and affinity estimates. Computational methods have significantly advanced individual stages of this pipeline, including structure prediction, generative design, and property prediction, yet these tools are typically developed in isolation. This disconnected landscape introduces significant decision friction: a structurally plausible pose does not guarantee strong binding, and a top-ranked sequence does not ensure biological function. Consequently, scientists are left to manually reconcile disjointed predictions, gauge model confidence, and prioritize candidates for costly experimental validation. To address this fragmentation, we introduce AnewDDE, an integrated Drug Discovery Engine that combines predictive foundation models with generative design and a scientific-domain post-trained language model.

AnewDDE operates as an agentic, closed-loop computational engine. Its structural foundation is anchored by AnewFold, which delivers reliable hypotheses for challenging targets, including anti-body–antigen complexes and molecular glues, and supports pocket identification and conformational analysis through AnewSampling [1]. AnewDesign then translates these insights into targeted candidates to reduce the experimental search burden. For quantitative evaluation, AnewAffinity combines near-FEP accuracy [2, 3] with the computational speed required for large-scale, high-throughput screening and for prioritizing promising transformations, and each prediction is accompanied by an FEP-like estimate of its own reliability. Finally, AnewMind, a post-trained, hundred-billion-parameter large language model (LLM) dedicated to scientific reasoning, bridges the gap between predictive outputs and expert decision-making, the very friction created by today’s fragmented workflows. It is evaluated on PharmBench, a benchmark designed by internal pharmaceutical R&D experts to assess long-horizon decision-making in drug discovery.

This report details the integrated components of AnewDDE in a progressive order:

- **AnewFold.** AnewFold is a general-purpose biomolecular structure prediction model covering antibody–antigen complexes, protein–ligand interactions, blind pocket identification, and molecular-glue ternary complexes. It delivers leading or competitive performance on recent, similarity-filtered benchmarks, achieving success rates of 60.8-76.2% for antibody structure prediction, 77.4% on the FoldBench protein–ligand benchmark, and 71.8% for post-cutoff molecular-glue complexes. AnewFold also captures cryptic pockets and ligand-induced conformational changes, while AnewSampling enables prospective validation of predicted interaction hotspots [1].
- **AnewAffinity.** We combine the rigorous physics of free-energy perturbation (FEP) with a distribution-predictive AI framework for fast, accurate, and scalable binding-affinity prediction. Beyond a single affinity estimate, AnewAffinity predicts the full free-energy distribution across FEP windows, enabling rigorous uncertainty estimation and sampling assessment. It yields relative binding affinities in *∼*1.5 seconds per ligand on a single GPU and achieves stronger correlation with experiment than Boltz-2 [4] (Δ*G* Pearson *R*^2^: 0.553 vs. 0.486; Spearman *ρ*: 0.724 vs. 0.620).
- **AnewDesign.** AnewDesign is an agentic lab-in-the-loop workflow that accelerates therapeutic biologics discovery by integrating generative AI models with wet-lab experimental feedback. In a proof-of-concept study targeting an internal protein target, the workflow combined *de novo* design with *in silico* affinity maturation and achieved 10.7% success rate in designing VHHs with single-digit nanomolar binding affinities from a panel of only 150 clones.
- **AnewMind.** We conduct full-parameter post-training at the hundred-billion-parameter scale and introduce PharmBench to evaluate long-horizon decision-making in drug discovery. The resulting AnewMind Preview matches or surpasses frontier LLMs in both ADMET prediction and rigorous pharmaceutical reasoning.

Looking ahead, combining the frontier-level performance of these proprietary tools with advanced scientific reasoning lays the foundation for a fully agentic scientific workflow, in which the LLM can seamlessly invoke individual modules on demand, integrate empirical feedback from each design– measure–learn cycle, and actively accelerate the translation of molecular insights into high-quality drug candidates.

## 2 Structure Modelling with AnewFold

### 2.1 Structure Prediction Across Biomolecular Interfaces

AnewFold is designed to model biomolecular recognition across multiple levels of structural complexity, from pairwise macromolecular interfaces to ligand-induced pockets and cooperative ternary complexes. We evaluate this capability in four complementary settings: antibody–antigen docking under stringent homology exclusion, protein–ligand co-folding across controlled novelty regimes, blind identification of constitutive and cryptic pockets, and molecular-glue ternary complex prediction. Together, these evaluations test whether AnewFold can recover not only globally plausible structures, but also the local interfaces, ligand poses, and conformational responses that determine molecular function. In all four settings, AnewFold consistently outperforms established co-folding baselines, with the largest gains on low-similarity targets and on interaction modes that are underrepresented in training data [5–9].

#### 2.1.1 Antibody-antigen Structure Prediction

Antibody–antigen interface prediction is one of the most demanding tasks in biomolecular co-folding. The binding surface is concentrated on the flexible complementarity-determining region (CDR) loops, which vary extensively in both sequence and conformation, and antigen epitopes are equally heterogeneous, leaving little homologous signal from which to infer the binding mode. We evaluated AnewFold on three public benchmark collections: AF3-AB, the antibody set used in the AlphaFold 3 (AF3) evaluation (71 entries, 65 clusters) [5, 10]; PXMeter-AB, which emphasizes therapeutically relevant antibodies (516 entries, 376 clusters) [11]; and FoldBench-AB, the low-homology antibody subset of FoldBench curated from recently released PDB structures (113 entries, 172 interfaces) [12].

AnewFold achieves top-1 success rates of 62.3% on AF3-AB, 60.8% on PXMeter-AB, and 76.2% on FoldBench-AB across five seeds (Fig. 2a, Table S1), surpassing Protenix-v2 by 8.8, 11.1, and 11.2 percentage points, respectively [8]. Its performance is comparable to that of the recently released IsoDDE model and ahead of AF3, Boltz-1, OpenFold3-preview2 (OF3p2), and OpenDDE [5, 9, 13, 14]. This performance is maintained across all evaluated quality thresholds (DockQ *>* 0.23, *>* 0.49, and *>* 0.8; Fig. 2a).

**Figure 1.**
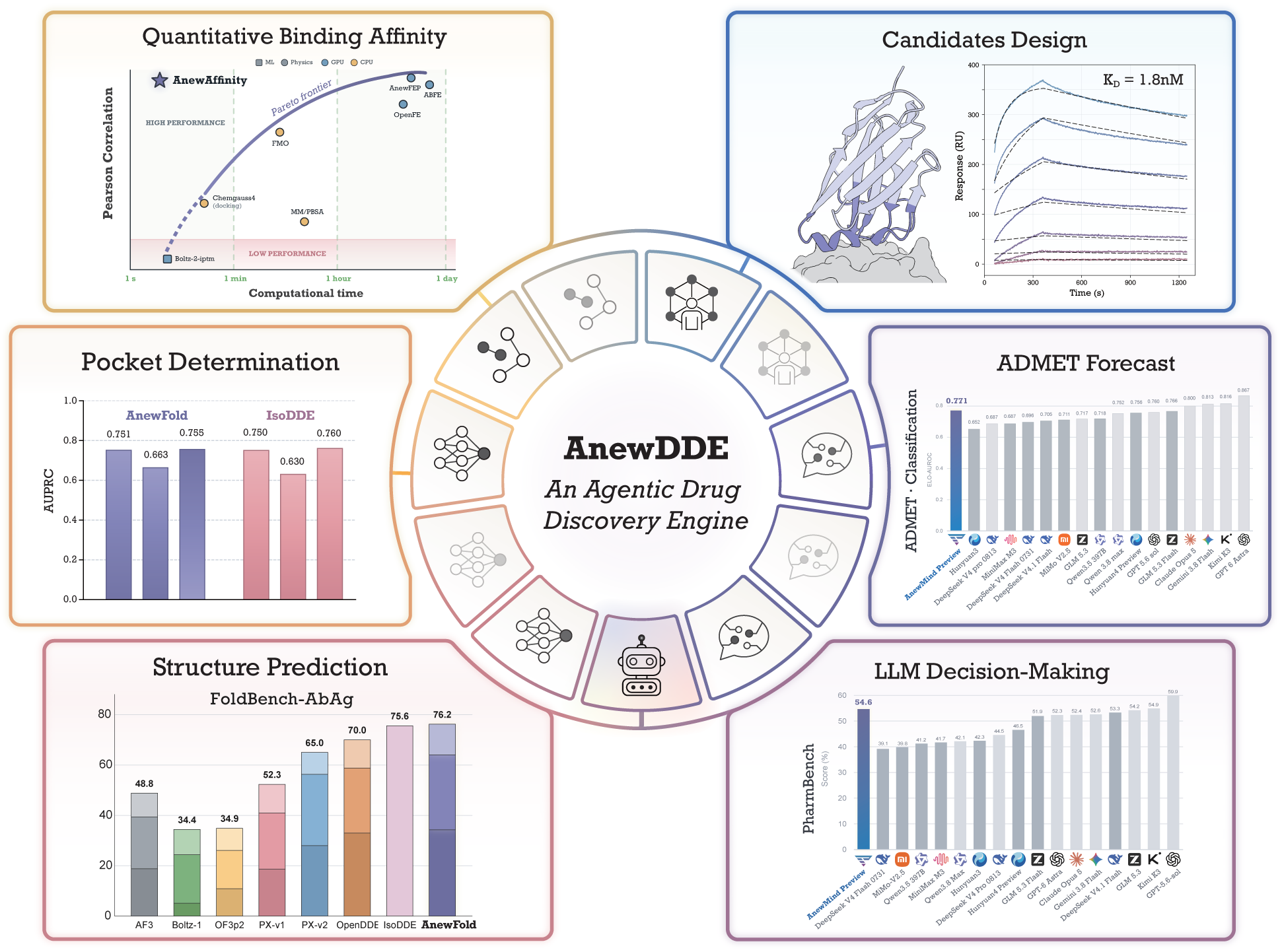
An agentic, closed-loop overview of AnewDDE and its key capabilities. Surrounding panels highlight representative evaluations across core capabilities. *Structure prediction*: AnewFold demonstrates high accuracy, outperforming several recent predictors. *Pocket determination*: robust pocket identification performance comparable to competitive baselines. *Quantitative binding affinity*: AnewAffinity balances prediction accuracy and computational speed, achieving high performance within a favorable time regime. *Candidate design*: successful *de novo* design yields high-affinity binders validated by experimental assays. *LLM post-train*: AnewMind Preview excels in ADMET property prediction and drug discovery long-horizon reasoning and decision making benchmarks.

**Figure 2.**
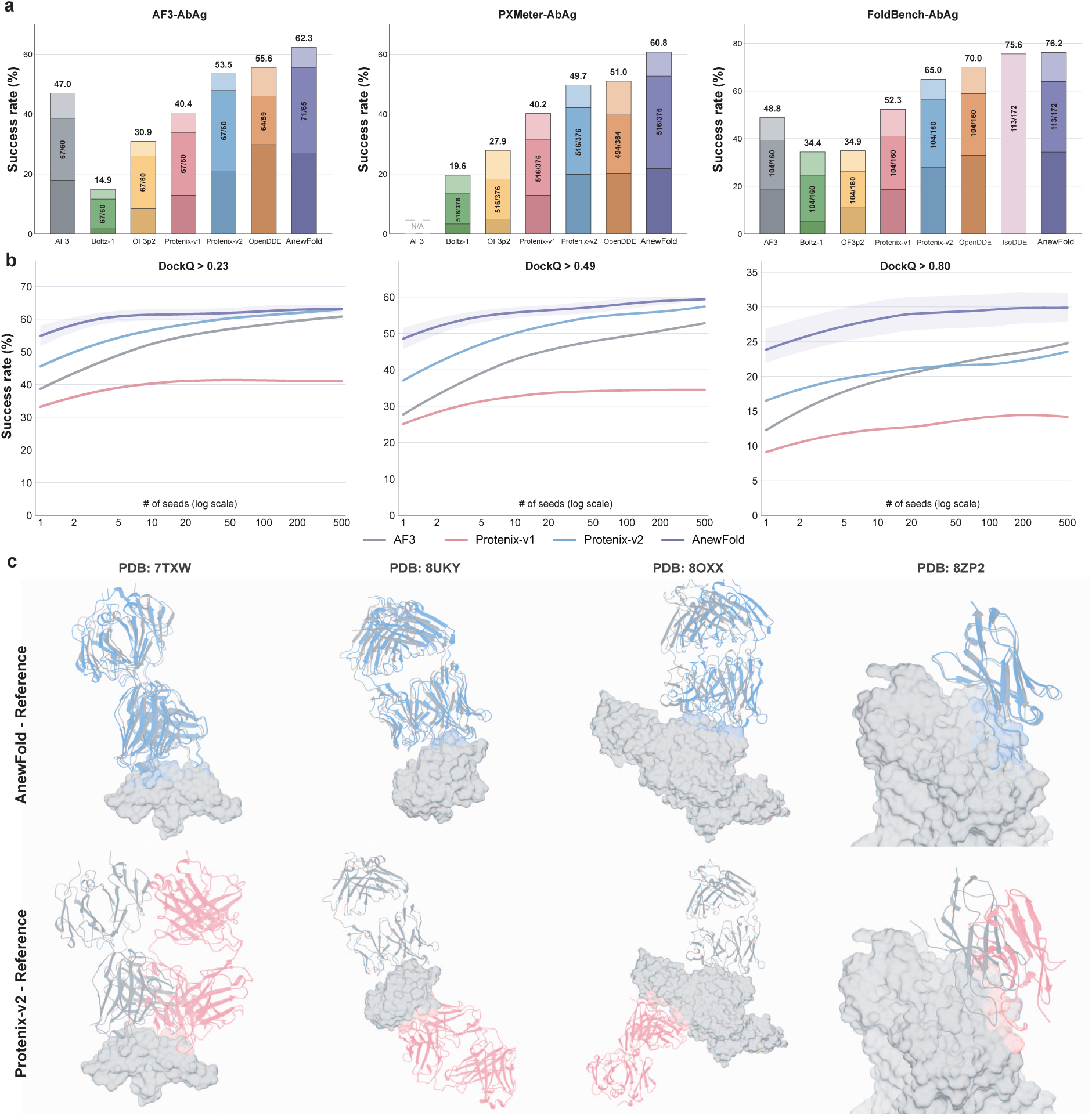
Antibody-antigen interface prediction. **(a)** Top-1 success rates across five seeds on three benchmark collections, AF3-AB, PXMeter-AB, and FoldBench-AB, with stacked bars denoting the DockQ *>* 0.8, 0.49 *<* DockQ *≤* 0.8, and 0.23 *<* DockQ *≤* 0.49 quality tiers. AnewFold achieves the highest success rate on all three collections: 62.3% on AF3-AB, 60.8% on PXMeter-AB, and 76.2% on FoldBench-AB. OpenDDE results are based only on targets with available predictions, including AF3-AB (64 entries/59 clusters), PXMeter-AB (494 entries/364 clusters) and FoldBench-AB (104 entries/160 interfaces). All AF3 results are taken from previously reported evaluations. **(b)** DockQ success rate as a function of the number of sampled seeds (1–500, log scale) at the DockQ *>* 0.23, *>* 0.49, and *>* 0.8 thresholds on AF3-AB (71 entities/65 clusters); shaded bands indicate bootstrap standard deviation. AnewFold leads at every seed budget and every quality threshold. **(c)** Representative predictions on four recent antibody–antigen targets (PDB 7TXW, 8UKY, 8OXX, 8ZP2). Top row: AnewFold predictions (blue) overlaid on the experimental reference (gray; antigen shown as surface); bottom row: Protenix-v2 predictions (pink). AnewFold places the antibody onto the correct epitope in the correct orientation, whereas Protenix-v2 either docks to a wrong antigen surface or binds in a wrong orientation.

AnewFold maintains its advantage as the number of sampled seeds increases on AF3-AB. From 1 to 500 seeds, it achieves higher success rates than AF3, Protenix-v1, and Protenix-v2 at all three DockQ thresholds (Fig. 2b). With a single seed, AnewFold already outperforms Protenix-v1 sampled with 500 seeds, and with tens to hundreds of seeds it matches AF3 and Protenix-v2, indicating substantially higher prediction accuracy per sample. The advantage also persists at DockQ *>* 0.8, showing that AnewFold improves high-accuracy interface geometry rather than merely raising the fraction of predictions above the minimum success threshold.

Qualitative examples from recent structures are consistent with the benchmark results (Fig. 2c). We highlight four held-out targets that probe generalization in the near absence of homologous structural priors, a regime in which Protenix-v2 consistently fails through interface inversion or epitope mislocalization. On the malaria transmission-blocking vaccine target Pfs25 bound to the neutralizing antibody 1G2 (PDB 7TXW) [15], AnewFold achieved near-native accuracy with a DockQ of 0.885 versus 0.028 for Protenix-v2, under sparse training coverage (81% CDR identity, 44% epitope similarity). For the pro-apoptotic effector BAK in complex with an inhibitory antibody (PDB 8UKY) [16], AnewFold mapped a novel, previously unobserved epitope to reach a DockQ of 0.614, whereas Protenix-v2 docked the antibody onto the opposing face of the protein (DockQ 0.031); the closest training exemplars show 77% CDR identity and only 7% epitope similarity. In the complex of transglutaminase 3 (TGM3) with a patient-derived Fab (PDB 8OXX) [17], AnewFold reconstructed the conformational epitope with a DockQ of 0.490 compared with 0.004 for Protenix-v2 (closest training CDR identity 61%, epitope similarity 8%). Finally, for the human noradrenaline transporter (NET) bound to a conformation-locking nanobody (PDB 8ZP2) [18], a membrane-protein system with no prior nanobody entries in the training data, AnewFold reached a DockQ of 0.894 compared with 0.103 for Protenix-v2 (70% CDR identity, 4% epitope similarity), extending the same generalization across membrane-protein topology. Such errors are not always fully captured by aggregate interface metrics, yet they directly affect the biological interpretation of a predicted complex. Together with the benchmark and inference-time scaling results, these examples support AnewFold’s ability to generalize to unseen antibody–antigen recognition modes under stringent homology exclusion.

These antibody–antigen results show that AnewFold can model diverse protein–protein interfaces without relying on close sequence or epitope homologs. We next evaluated its generalization to protein–ligand complexes, which additionally requires correct pocket identification, ligand placement, and local interaction geometry.

#### 2.1.2 Protein-Ligand Structure Prediction

A central question for co-folding models is whether ligand poses are recovered from genuine physical modeling rather than from templates resembling the training data. We first evaluated AnewFold on FoldBench Protein–Ligand, whose complexes were deposited after January 13, 2023, and were filtered for similarity to structures before the date [12]. Under the top-1 joint success criterion (pocket-aligned ligand RMSD below 2.0 Å together with LDDT-PLI above 0.8), AnewFold achieves 77.4%, exceeding IsoDDE (76.0%) [9], Protenix-v2 (64.0%) [8], Protenix-v1 (62.8%) [7], AF3 (64.9%) [5], OpenDDE (60.1%) [20] and OF3p2 (44.5%)(Fig. 3a).

**Figure 3.**
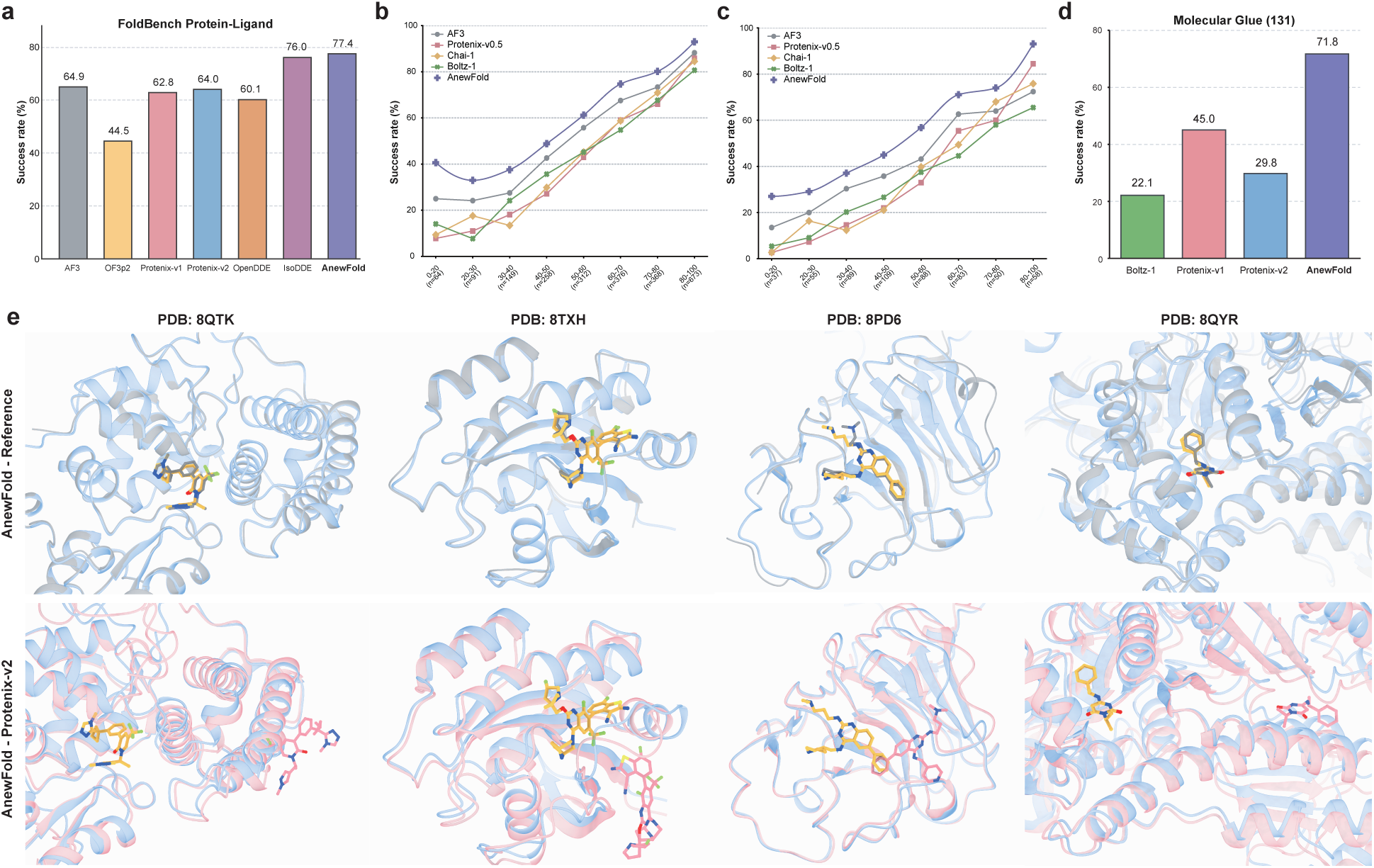
Protein-ligand prediction under in-distribution, similarity-stratified, and out-of-distribution settings. **(a)** Top-1 joint success rate on FoldBench Protein–Ligand, comprising complexes deposited after January 13, 2023, under stringent similarity filtering. AnewFold reaches 77.4%, compared with 76.0% for IsoDDE, 64.0% for Protenix-v2, 62.8% for Protenix-v1, 64.9% for AF3, and 44.5% for OF3p2. **(b, c)** RNP benchmark results. **(b):** Full dataset, **(c):** clustered and filtered to remove 124 prevalent ligands, as in [19]. Success rate is plotted as a function of the joint pocket–ligand shape similarity between each test complex and its nearest training-set neighbor, binned from [0.0, 0.2] to [0.8, 1.0]. AnewFold remains the top-ranked model across the entire similarity range, including the lowest similarity bins where the baselines degrade most strongly. **(d)** Top-1 joint success rate on our newly curated molecular-glue benchmark (131 ternary complexes deposited after September 30, 2021): Boltz-1 22.1%, Protenix-v1 45.0%, Protenix-v2 29.8%, and AnewFold 71.8%. **(e)** Representative out-of-distribution cases. Top row: AnewFold predictions (light blue) overlaid on the experimental reference (gray); bottom row: the same predictions compared with Protenix-v2 (pink), with ligands shown as yellow sticks. From left to right: the Cbl-b allosteric pocket (PDB 8QTK), the KRAS G12D Switch-II cryptic pocket (PDB 8TXH), the TRIM58 PRY-SPRY substrate-recognition pocket (PDB 8PD6), and the mavacamten–*β*-cardiac myosin interface allosteric pocket (PDB 8QYR).

To evaluate protein–ligand generalization, we used the Run N’ Poses (RNP) benchmark, which stratifies each test complex by its joint pocket–ligand shape similarity to the nearest training-set example [19]. Across all similarity bins (Fig. 3b,c), AnewFold achieves higher success rates than AF3 [5], Chai-1 [21], Boltz-1 [13], and Protenix-v0.5 [6]. The advantage is most pronounced in the lowest-similarity bins, where baseline performance declines sharply while AnewFold remains comparatively stable. These results suggest that AnewFold is less dependent on close pocket–ligand analogs in the training set.

Molecular glues are particularly challenging to design because their activity depends on promoting or stabilizing specific protein–protein interactions rather than binding to a single protein alone. We extended the MGTbind structural collection [22] with PDB depositions, yielding 131 complexes deposited after Sep. 30, 2021. On this after-cutoff set, AnewFold reaches a 71.8% top-1 joint success rate (DockQ *>* 0.23, pocket-aligned ligand RMSD *<* 2.0 Å, and LDDT-PLI *>* 0.8), versus 45.0% for Protenix-v1, 29.8% for Protenix-v2, and 22.1% for Boltz-1 [13] (Fig. 3d and Fig. S1). The large gain over all baselines on these three-component systems shows that AnewFold’s advantage extends to molecular glues and their characteristic ligand-induced protein–protein interfaces, which are not represented in most conventional docking benchmarks.

Benchmark-level success rates do not reveal whether a model places the ligand in the biologically correct pocket, particularly for allosteric and cryptic sites. Four recent structures illustrate AnewFold’s performance in these settings (Figure 3e).

##### Cbl-b allosteric pocket (PDB 8QTK)

The ligand binds an induced-fit allosteric pocket, driving local side-chain rearrangements that stabilize the autoinhibited conformation of Cbl-b [23]. Despite a nearest-neighbor pocket–ligand similarity of only 5.4%, AnewFold reproduces the experimental binding site and ligand pose, whereas Protenix-v2 fails to localize the pocket.

##### KRAS G12D Switch-II pocket (PDB 8TXH)

The ligand occupies the Switch-II cryptic pocket formed through local conformational rearrangement [24–26]. AnewFold places the ligand within this dynamically induced pocket with the correct orientation, while Protenix-v2 predicts a pose on a pre-existing surface region.

##### TRIM58 PRY-SPRY domain (PDB 8PD6)

TRIM58 is a RING-type E3 ligase expressed in erythroid precursor cells [27]; the ligand engages its PRY-SPRY substrate-recognition domain, a recruitment interface whose nearest-neighbor ligand-and-pocket shape similarity to the training set is below 14%. AnewFold recovers the experimental pose in the substrate-recognition pocket, whereas Protenix-v2 assigns the ligand to an incorrect site.

##### Mavacamten-*β*-cardiac myosin (PDB 8QYR)

Mavacamten occupies the inter-domain allosteric pocket between the myosin motor domain and the lever arm, locking the motor in its pre-powerstroke state [28]. At 27% nearest-neighbor similarity, AnewFold correctly identifies this pocket, whereas Protenix-v2 places the ligand in the orthosteric ADP-binding site.

Across these examples, AnewFold recovers allosteric, cryptic, and inter-domain binding sites at nearest-neighbor similarities ranging from 5.4% to 27%. Protenix-v2 instead favors a pre-existing or orthosteric pocket, or fails to identify the binding site. Together with the similarity-stratified RNP results and the post-cutoff molecular-glue benchmark, these cases indicate that AnewFold generalizes to protein–ligand interactions that differ substantially from the training data. These results address ligand-conditioned co-folding, in which the query ligand is provided to the model. We next consider the more general problem of identifying binding sites directly from protein sequences, including cryptic pockets that become accessible only after conformational rearrangement [25, 26, 29].

### 2.2 Pocket Identification

The structure prediction described above presupposes a given ligand. Pocket identification answers a more fundamental question: in the absence of any ligand information, can the model directly identify all potential binding sites on a protein? Unlike ligand-conditioned structure prediction, blind pocket identification asks whether a model can locate potential binding sites directly from the protein sequence. This capability, also reported for IsoDDE [9], is relevant to discovering druggable sites on novel targets and identifying alternative mechanisms such as allosteric regulation. We therefore evaluated AnewFold in a ligand-free pocket identification setting, covering both constitutive and cryptic pockets (see Supplementary Information for details).

We evaluated binding-pocket residue prediction using the area under the precision–recall curve (AUPRC), which is well suited to the strong class imbalance between pocket and non-pocket residues. AnewFold achieves an overall AUPRC of 0.751, with 0.755 on non-cryptic pockets and 0.663 on cryptic pockets (Fig. 4a). The lower AUPRC for cryptic pockets reflects the additional difficulty of detecting binding sites that are only partially formed or inaccessible in the unliganded structure. On its own evaluation set, IsoDDE reports similar AUPRC values: 0.750 overall, 0.760 for non-cryptic pockets, and 0.630 for cryptic pockets [9]. These values are not directly comparable, because the two models were assessed on independent held-out collections with different target distributions, pocket annotations, and cryptic-site definitions; they nevertheless indicate comparable accuracy under their respective protocols.

**Figure 4.**
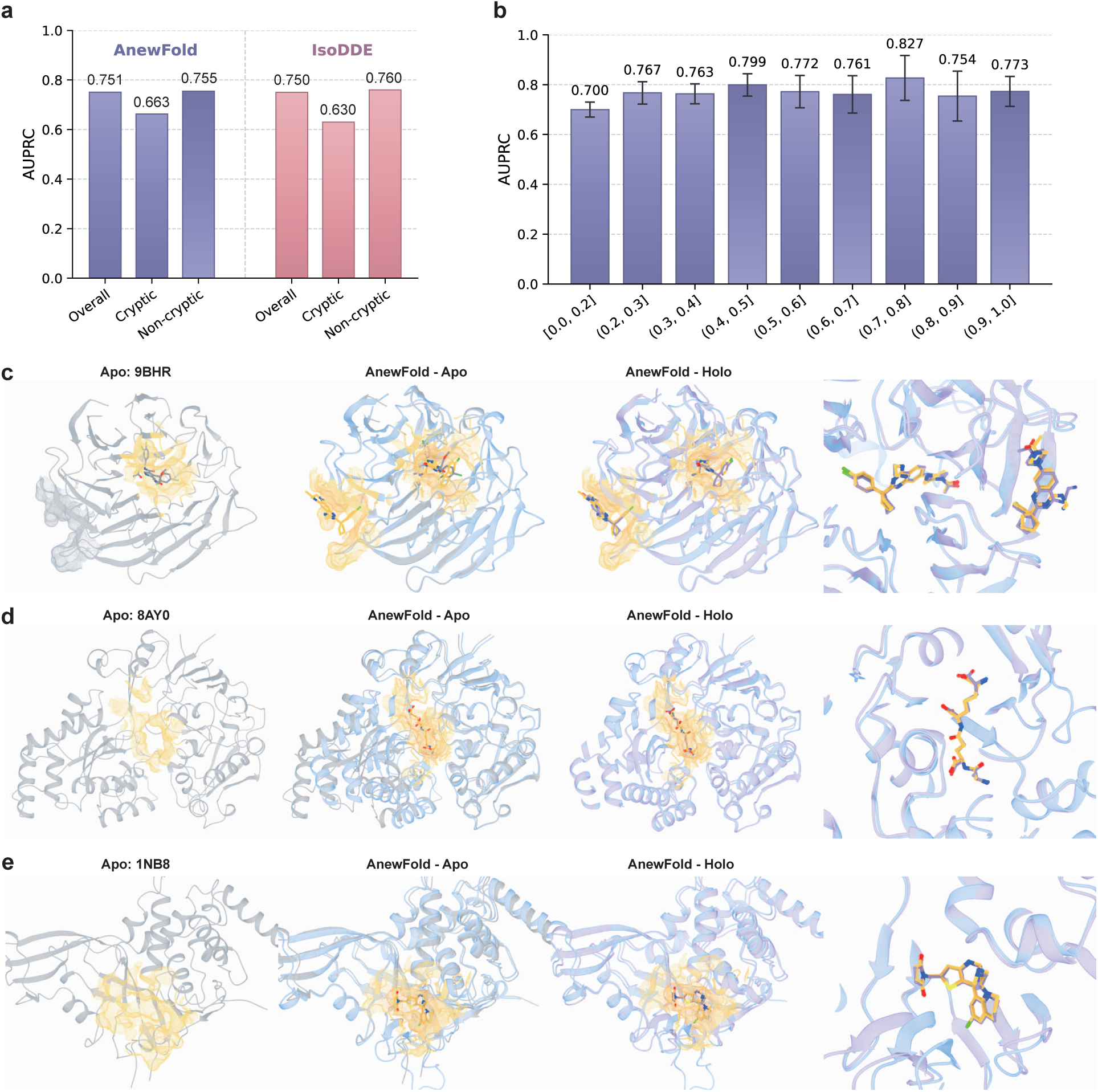
Pocket prediction and apo-to-holo conformational modeling by AnewFold. **(a)** Pocket prediction measured by AUPRC. AnewFold achieves 0.751 overall, 0.663 on cryptic pockets, and 0.755 on non-cryptic pockets; the corresponding values for IsoDDE are 0.750, 0.630, and 0.760. Because the two models were evaluated on different held-out sets, their performance are shown for reference rather than comparison. **(b)** AnewFold AUPRC stratified by sequence identity between each test target and its nearest neighbor in the training set. Performance remains stable across similarity bins. Error bars denote bootstrap standard errors. **(c-e)** Representative apo-to-holo conformation predictions. From left to right in each row: the experimental apo structure, the AnewFold prediction conditioned on the apo structure, and the prediction overlaid with the experimental holo structure, with a close-up of the pocket and ligand. **(c)** An allosteric binder induces localized pocket formation in apo 9BHR (holo 8OGC, C*α* RMSD 0.9 Å). **(d)** Ligand binding coupled to an open–close domain motion in apo 8AY0 (holo 8AZB, C*α* RMSD 4.0 Å). **(e)** A local closed-to-open pocket motion in apo 1NB8 (holo 8D4Z, C*α* RMSD 1.1 Å). Predicted pockets are highlighted as yellow surfaces, predicted proteins are shown in light blue/purple and reference structures in gray/pink, with ligands as sticks.

To test whether this performance depends on close homologs, we stratified the evaluation set by the sequence identity between each test target and its nearest training-set neighbor (Fig. 4b). AUPRC remains within 0.700–0.827 across all bins and retains a value of 0.700 for targets sharing at most 20% sequence identity with any training protein, indicating that pocket identification is driven by local structural and physicochemical features of the binding site rather than by memorization of close homologs.

The examples in Fig. 4c–e examine whether AnewFold can recover holo-like conformations from apo inputs. For the WD40 *β*-propeller domain of DCAF1 (apo 9BHR, holo 8OGC) [30], AnewFold reproduces the lateral allosteric cleft between blades 1 and 2 while retaining the central orthosteric channel, with an overall C*α* RMSD of 0.9 Å and pocket- and ligand-alignment RMSDs of 0.85 and 1.12 Å. For the *Bacillus subtilis* substrate-binding protein DppE (apo 8AY0, holo 8AZB) [31], ligand binding closes the two domains in a Venus-flytrap motion around the murein-tripeptide ligand; AnewFold recovers both the closed conformation and the inter-domain binding groove (pocket RMSD 0.42 Å, ligand RMSD 0.85 Å). For USP7 (apo 1NB8, holo 8D4Z) [32], the cryptic allosteric pocket is collapsed in the apo structure, where the catalytic triad sits unaligned in an inactive, open state. AnewFold recovers the ligand-opened pocket (pocket- and ligand-alignment RMSDs of 1.46 Å) together with the inhibitor-bound conformation, in which the allosteric ligand wedges the pocket open and locks the switching loop in its inactive, open state, blocking the closure required for catalytic activation.

Blind pocket identification tests binding-site recognition without ligand information. A stricter test is whether this capability remains coupled to structure prediction in a multicomponent system, where a small molecule creates a new protein–protein interface rather than occupying an isolated pocket. We therefore analyzed the KAT2A–Compound 4(PDB CCD: A1C5C)–CRBN(Cereblon) molecular-glue complex using experimentally validated hotspot mutations to determine whether the conformational ensemble preserves the contacts required for glue-mediated recruitment [22, 33].

### 2.3 A Representative Case Study of a Molecular-glue Ternary Complex

Molecular glues are small molecules that induce or stabilize protein–protein interactions, thereby modulating protein function. Predicting their mode of action therefore requires accurate modeling of both the ligand pose and the induced protein–protein interface within a ternary complex [22]. On the 131-complex molecular-glue benchmark described above, AnewFold shows a substantial improvement over the evaluated baselines (Boltz-1, Protenix-v1, Protenix-v2) shown in Fig. 3d and Fig. S1. To examine whether this improvement extends to mechanistically important contacts, we further analyzed a recently reported system in which A1C5C recruits the histone acetyltransferase KAT2A to the CUL4–DDB1–CRBN ubiquitin ligase by stabilizing the KAT2A–CRBN interaction [33]. The experimental structure shows that the ternary complex is stabilized by three cooperative contact patches: a compound interface, in which A1C5C wraps around KAT2A Y200; a helix interface, in which the termini of two KAT2A *α*-helices contact the zinc-coordinating region of the CRBN CTD; and a loop interface, in which the F150-containing loop of the CRBN NTD contacts KAT2A residues adjacent to its binuclear zinc center. Importantly, both protein interfaces are functionally required: combined mutation of the helix residues (K289A/V285A/R277A) or of the loop residues (F109E/F202E/F183A/R206A) abolishes KAT2A recruitment in cells [33]. This system therefore provides a stringent, mutagenesis-validated test of whether a co-folding model recovers the correct ternary geometry rather than merely docking the compound into a single pocket.

AnewFold predicts the ternary complex directly from the KAT2A and CRBN sequences with A1C5C, without using the experimental structure as a template (Fig. 5a). Using AnewSampling [1], a cofolding-like architecture designed for equilibrium conformational sampling, we generated an ensemble of 500 conformers. All three experimentally identified contact patches were recovered across the sampled ensemble (Fig. 5b,c). The loop interface is the most robustly captured: the CRBN-F150/KAT2A contacts and the KAT2A hotspot residues are preserved in 97.8–99.6% of conformers (F109 98.4%, F183 97.8%, R206 99.0%, CRBN-F150 99.6%), consistent with the extensive hydrophobic footprint of this interface. The helix interface is recovered in an asymmetric pattern, with R277 contacting CRBN in 88.0% of conformers, while K289 and V285 appear less frequently (18.8% and 8.0%); the compound-anchoring contact Y200 is present in 35.8% of conformers and F202 in 46.6%. The lower frequencies for these residues are compatible with their roles as conformation-dependent contacts rather than persistent anchors, and the ensemble as a whole retains at least one helix-interface contact in the large majority of predictions.

**Figure 5.**
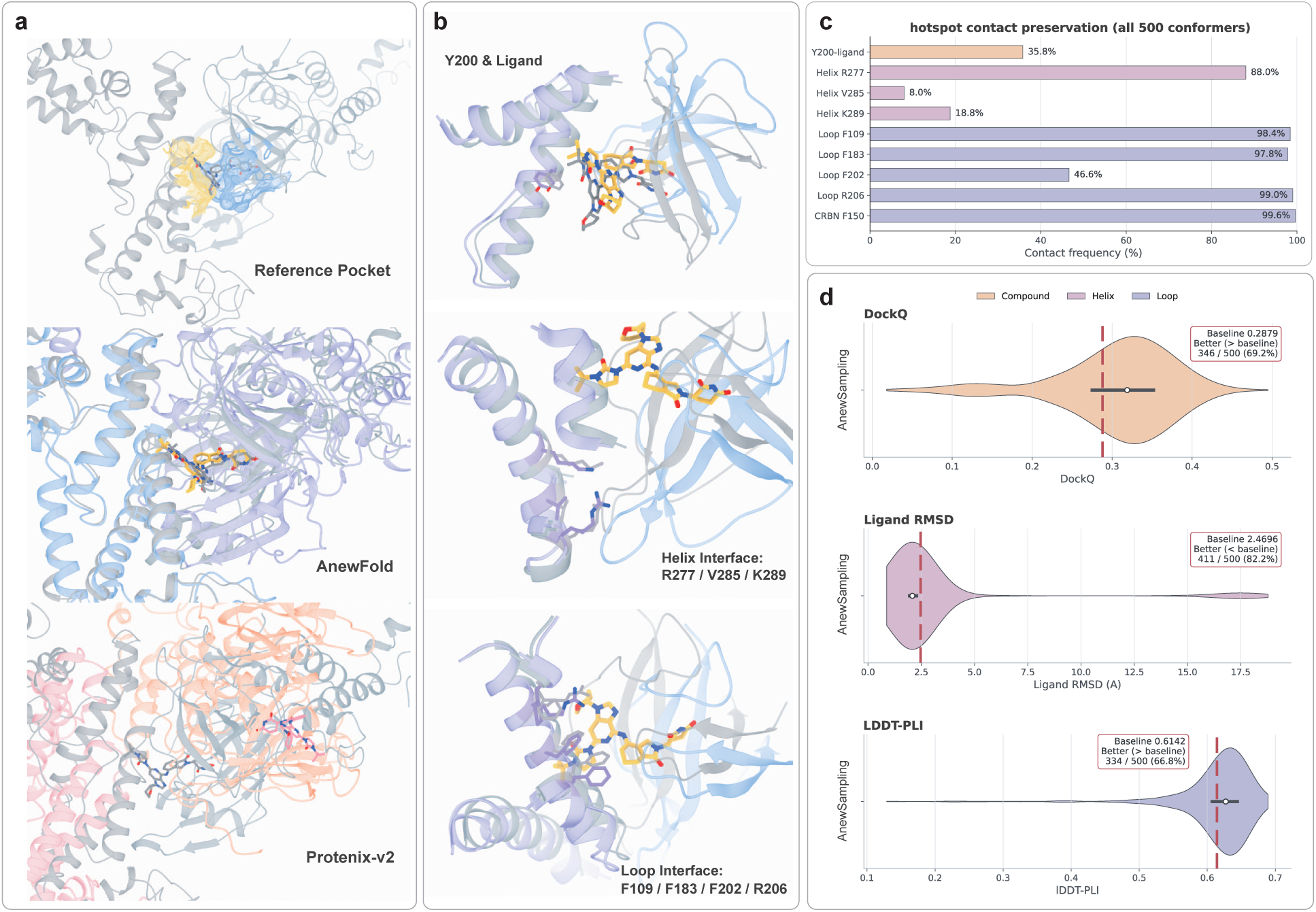
Molecular-glue ternary complex prediction for the KAT2A-A1C5C-CRBN system. **(a)** Predicted ternary complexes compared with the experimental reference. Top: experimental reference structure (gray); middle: representative AnewFold prediction; bottom: Protenix-v2 prediction. KAT2A is shown in light blue/purple, CRBN in light red/orange, and A1C5C as yellow sticks. **(b)** Close-up views of the three contact patches that stabilize the ternary assembly: the compound interface, where A1C5C wraps around KAT2A Y200; the helix interface, where the termini of two KAT2A *α*-helices contact the zinc-coordinating region of the CRBN CTD (KAT2A R277/V285/K289); and the loop interface, where the F150 loop of the CRBN NTD contacts KAT2A residues near the binuclear zinc center (KAT2A F109/F183/F202/R206, and CRBN-F150). **(c)** Frequency of the experimentally validated hotspot contacts across all 500 predicted conformers. Loop-interface contacts are the most stably preserved (CRBN-F150 99.6%, KAT2A F109 98.4%, F183 97.8%, R206 99.0%), followed by helix-interface R277 (88.0%); the compound-contact Y200 (35.8%) and the remaining helix/loop residues are recovered less frequently (KAT2A K289 18.8%, V285 8.0%, F202 46.6%). **(d)** Distribution of the 500 AnewSampling conformers (red dashed line: Protenix-v2 baseline) for the KAT2A/CRBN interface DockQ (baseline 0.2879; 346/500, 69.2% above baseline), ligand RMSD in Å (baseline 2.4696 Å; 411/500, 82.2% at or below baseline), and LDDT-PLI (baseline 0.6142; 334/500, 66.8% above baseline).

We further quantified ensemble-level accuracy on three orthogonal metrics (Fig. 5d). For the induced KAT2A-CRBN interface, 346 of 500 conformers (69.2%) exceed the baseline DockQ of 0.2879. For the ligand, 411 of 500 conformers (82.2%) achieve a ligand RMSD no greater than the 2.4696 Å baseline, with the distribution concentrated well below 2.5 Å. On the per-residue protein–ligand interaction score LDDT-PLI, 334 of 500 conformers (66.8%) surpass the baseline value of 0.6142.

Taken together, these results connect predicted geometry to experimental causality. The contacts AnewFold preserves most frequently across the 500-conformer ensemble— the CRBN-F150 loop against KAT2A F109/F183/R206 and the KAT2A helix R277 against the CRBN zinc-coordinating region—are the patches whose combined mutagenesis eliminates recruitment [33]. Agreement with both the three-interface structural model and the two loss-of-recruitment mutant sets indicates that AnewFold captures the cooperative, glue-induced recognition mechanism of a previously unseen molecular-glue ternary complex, rather than predicting a plausible but non-functional binding pose.

Across antibody–antigen complexes, protein–ligand systems, blind pocket identification, and molecular-glue ternary assemblies, AnewFold exhibits a consistent pattern: its advantage grows as the task moves away from close training analogs and toward interaction modes governed by flexible loops, cryptic cavities, induced fit, or cooperative interfaces. The model not only raises benchmark success rates, but also recovers mechanistically specific features: the correct antibody epitope and orientation, low-similarity ligand poses, apo-to-holo pocket opening, and mutagenesis-supported molecular-glue hotspots. These results support a unified view of AnewFold as a general biomolecular interaction model that couples global assembly with local chemical and conformational accuracy, providing a stronger structural basis for downstream therapeutic discovery than template-centered prediction alone.

## 3 Quantitative Binding with AnewAffinity

### 3.1 Fast, Accurate, and Inspectable Affinity Prediction

Quantitative affinity prediction is what turns a structural hypothesis into a design decision: the complexes and poses from AnewFold define candidate transformations, while their relative free energies provide a central quantitative criterion for prioritizing which ligands to synthesize. AnewAffinity predicts relative protein–ligand binding free energies with near-FEP accuracy at seconds-per-edge cost, screening entire perturbation networks within a single design cycle, and every prediction is accompanied by an FEP-like reliability estimate, supported by phase- and window-resolved evidence that quantifies the confidence in each individual prediction rather than reporting only an aggregate score. It thereby resolves a dilemma that existing methods leave open: physics-based FEP is accurate and diagnostically rich but too slow for large-scale screening, whereas fast learned scorers cover the design space yet return a single opaque number, with no built-in mechanism to assess the reliability or confidence of an individual prediction or to diagnose where it fails. We first make this trade-off explicit through the strengths and limits of physical FEP, and then describe how AnewAffinity retains its inspectable evidence at learned-model speed.

Physics-based relative binding free-energy (RBFE) calculations set a high accuracy standard because they evaluate a thermodynamic cycle rather than mapping structure directly to affinity; their sampled work distributions provide both an estimate and transformation-specific evidence on overlap and convergence [2, 3]. The same sampling, however, limits throughput: each transformation requires molecular dynamics across many alchemical windows in both phases, so a network of tens or hundreds of edges expands into thousands of simulations. The cost also exceeds GPU time. When a prediction deviates from experiment, distinguishing sampling deficiencies—such as poor phase-space overlap, side-chain or ligand-pose transitions, water-network rearrangements, or suboptimal *λ* schedules—from force-field bias often requires an iterative, expert-driven debugging process involving repeated simulations and, when necessary, reparameterization [3, 34]. This process can be slower than medicinal-chemistry design cycles allow.

AnewAffinity closes this gap with near-kcal/mol retrospective error, stronger aggregate Δ*G* correlation, and better ligand ranking than the Boltz-2 evaluation described below, at an end-to-end cost of approximately 1.5 seconds per ligand-pair edge. Rather than a bare scalar, each estimate carries phase- and window-resolved, FEP-compatible evidence—directional work distributions, overlap information, and locally accumulated free energies of the kind a physicist inspects in an explicit simulation—so internal consistency can be checked and the specific window or phase behind a questionable result localized, as demonstrated for a held-out BACE transformation in Fig. 8 and Table 2.

**Table 1.** Retrospective accuracy on the eight-system JACS benchmark. Each metric is computed within a system and aggregated across systems with ligand-count weights. Lower pairwise ΔΔ*G* RMSE and higher Δ*G* correlations are better.

| Method | Ligands | $\Delta\Delta G$ RMSE (kcal/mol) | $R^2_{\text{Pearson}}$ | Spearman $\rho$ |
| --- | --- | --- | --- | --- |
| Boltz-2 | 199 | **1.016** | 0.486 | 0.620 |
| AnewAffinity | 199 | 1.057 | **0.553** | **0.724** |

**Table 2.** Agreement between AnewAffinity and reference FEP samples for the BACE transformation. Wasserstein-1 (W1) distance measures agreement between work distributions, and window Δ*G* MAE measures agreement between the resulting local free-energy estimates.

| Phase | Windows | Forward W1 ($RT$) | Backward W1 ($RT$) | $\Delta G$ MAE (kcal/mol) |
| --- | --- | --- | --- | --- |
| Protein | 23 | 0.246 | 0.246 | 0.133 |
| Water | 23 | 0.299 | 0.314 | 0.180 |

TYK2, JNK1, and p38 subset divided by ligand count—paired with a Pearson correlation of 0.76. The distinction matters operationally: per-edge latency governs how quickly a single new hypothesis can be checked during a design discussion, whereas amortized per-ligand cost governs the throughput available when an entire network is scored, and conflating the two would make head-to-head timing comparisons misleading.

### 3.2 Retrospective Accuracy Across Eight Protein Targets

We evaluated AnewAffinity on 199 ligands drawn from eight retrospective JACS systems spanning distinct target families and binding-site chemistries: BACE, CDK2, JNK1, MCL1, p38, PTP1B, thrombin, and TYK2. The series range from 11 to 42 ligands and cover both tight, narrow affinity ranges and wider multi-log-unit spreads, so the benchmark probes both numerical precision within a series and discrimination across chemically diverse targets. Three complementary metrics are used. Pairwise ΔΔ*G* RMSE measures numerical agreement for the specific transformations a team would act on, while squared Pearson correlation for Δ*G* and Spearman rank correlation measure whether affinity trends and ligand ordering are preserved—the difference between getting magnitudes right and getting the ordering right that drives prioritization. Each metric is computed within a system and aggregated across systems with ligand-count weights, so that systems with more measured compounds contribute proportionally and the summary cannot be dominated by one small series.

As summarized in Table 1, AnewAffinity achieves a pairwise ΔΔ*G* RMSE of 1.057 kcal/mol, a weighted *R*^2^ of 0.553, and a weighted Spearman *ρ* of 0.724. A local evaluation of the released Boltz-2 model [4] on the same 199 ligands gives 1.016 kcal/mol, 0.486, and 0.620, respectively.

Near-kcal/mol error is meaningful at the scale of medicinal-chemistry decisions: at room temperature, roughly 1.4 kcal/mol corresponds to a factor of ten in binding affinity. The two models show near-identical numerical RMSE, differing by only about 0.04 kcal/mol, whereas AnewAffinity attains a higher weighted Pearson *r*^2^ (0.553 vs. 0.486) and a clearly better ranking. Its advantage is therefore concentrated on candidate ordering rather than uniformly smaller error. More accurate physics-based approaches occupy a different, hour-scale operating regime, motivating the runtime comparison below.

Fig. 6 shows that performance varies across targets, with neither method dominating every system: on some series the two are close, while on others one is clearly ahead. Confidence in a prediction should therefore follow the target and chemical context rather than being inferred from a single aggregate score. This target-dependence is also where the per-transformation evidence of Section 3.4 becomes useful, because a benchmark-wide number cannot by itself say whether a particular edge lies in a well- or poorly-predicted region. On the ligand-count-weighted aggregate, however, AnewAffinity supplies the accuracy needed for prioritization while preserving stronger affinity trends and ligand ranking (*R*^2^ = 0.553, *ρ* = 0.724) across this heterogeneous target set.

**Figure 6.**
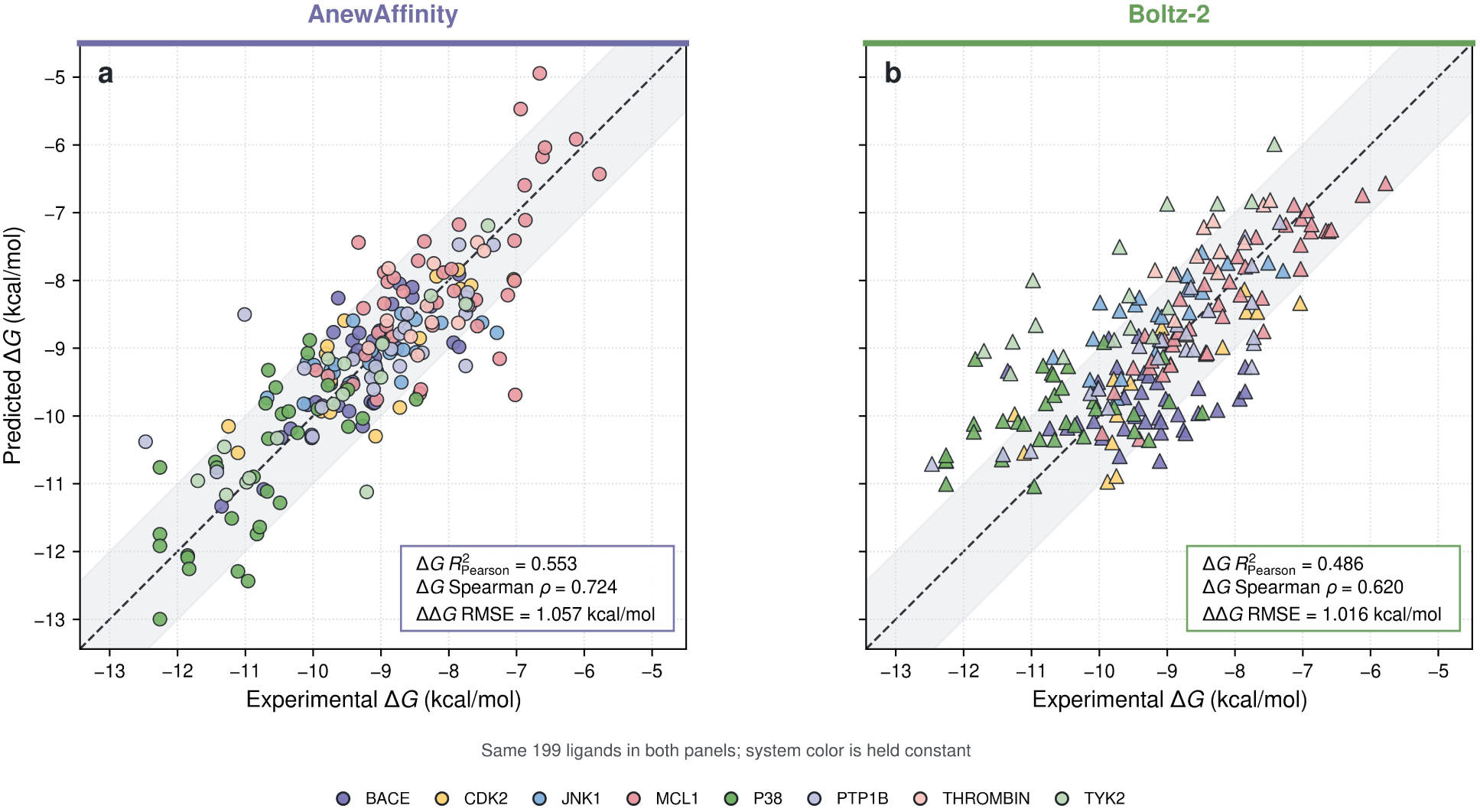
Δ*G* prediction across eight JACS systems. **(a)** AnewAffinity predictions, shown as circles. **(b)** Boltz-2 predictions, shown as triangles. Color identifies the protein system and is held constant between panels. Both methods are evaluated on the same 199 ligands with identical axis limits. Dashed lines indicate identity, and shaded bands indicate deviations within 1 kcal/mol. Insets report ligand-count-weighted squared Pearson correlation for Δ*G*, Spearman correlation for Δ*G*, and pairwise ΔΔ*G* RMSE.

### 3.3 Low-Latency Evaluation for Broader Design

Across GPU runs on multiple machines, AnewAffinity evaluates one ligand-pair edge in approximately 1.5 seconds end to end, including loading, inference, post-processing, and serialization. This per-edge latency is the marginal cost of adding a transformation and is distinct from the runtime coordinate in Fig. 7, which reports an amortized 3.4 seconds per ligand—the total GPU runtime over the CDK2, TYK2, JNK1, and p38 subset divided by ligand count—paired with a Pearson correlation of 0.76. The distinction matters operationally: per-edge latency governs how quickly a single new hypothesis can be checked during a design discussion, whereas amortized per-ligand cost governs the throughput available when an entire network is scored, and conflating the two would make head-to-head timing comparisons misleading.

**Figure 7.**
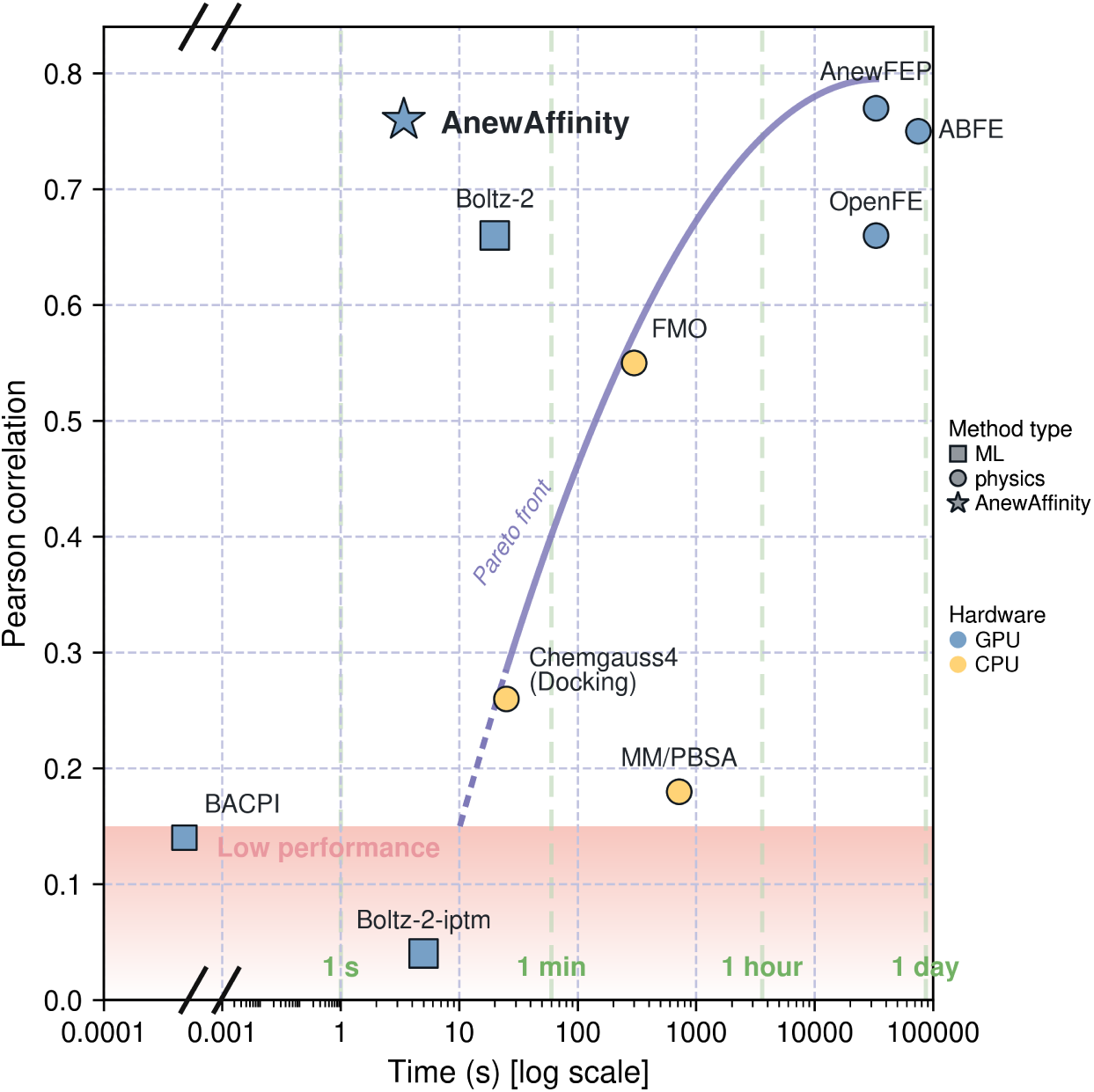
Runtime-accuracy landscape across representative computational approaches. Runtime is shown on a logarithmic axis and predictive accuracy by Pearson correlation. Fill denotes GPU or CPU hardware; squares denote machine-learning methods, circles denote physics-based methods, and the star highlights AnewAffinity. The shaded band marks Pearson correlations below 0.15, and vertical guides mark 1 second, 1 minute, 1 hour, and 1 day. The curve is an illustrative guide to the runtime–accuracy frontier traced by representative physics-based methods, rather than a fitted relationship. AnewAffinity uses the separate four-system measurement of an amortized 3.4 seconds per ligand (total GPU runtime divided by ligand count) with Pearson correlation 0.76; this differs from the 1.5-second per-edge latency reported in the text. AnewFEP is shown at 33,000 seconds and 0.77. Values span different datasets, hardware, and timing definitions and therefore provide a qualitative comparison rather than a controlled benchmark.

Fig. 7 compares runtime and predictive accuracy, reported as Pearson correlation *r*, across representative alternatives. Physics-based methods such as AnewFEP (*∼*33,000 seconds, *r* = 0.77), ABFE (*∼*75,000 seconds, *r* = 0.75) [35], and OpenFE (*∼*33,000 seconds, *r* = 0.66) [36], together with Boltz-2 at 20 seconds and *r* = 0.66 [4, 34], occupy more expensive operating regimes. IsoDDE reports a higher mean correlation (*r* = 0.85) on the separate FEP+ 4 benchmark, but no comparable runtime is available [9], so it cannot be placed on this landscape; hardware, sources, and the IsoDDE entry are reported in Supplementary Table S2. Because the points combine different datasets, hardware, and timing definitions, the figure is a qualitative operating-space map rather than a controlled speedup benchmark. In AnewDDE, second-scale evaluation allows affinity to guide candidate generation, network construction, and synthesis prioritization early in each design cycle.

### 3.4 Window-Resolved Free-Energy Evidence

The held-out BACE transformation CAT-24 *→* CAT-17e shows what interpretability means in practice. AnewAffinity does not return a single endpoint score: it evaluates the transformation separately in the protein and water phases, with phase-specific changes Δ*G*_P_ and Δ*G*_W_, and combines them through the thermodynamic cycle as ΔΔ*G* = Δ*G*_P_ *−* Δ*G*_W_. Within each phase it reports forward and backward work distributions at every alchemical window (23 windows per phase), the same quantities a physicist inspects in explicit FEP. At each window, the Bennett acceptance ratio (BAR) [37] combines the two directional work distributions—evaluated on the original, unreflected samples—to estimate the local free-energy difference; the window estimates are then accumulated over the windows and subtracted between phases. Weak or inconsistent forward/backward overlap at any window directly flags that transformation for further attention.

For CAT-24 *→* CAT-17e, AnewAffinity predicts ΔΔ*G* = +1.12 kcal/mol, close to the experimental +1.33 kcal/mol. Agreement is not confined to the final number. As summarized in Table 2, the predicted work distributions match reference FEP samples closely in both phases (mean W1 distance 0.246–0.314 *RT*), and the window-level Δ*G* errors are small (0.133 and 0.180 kcal/mol), so the estimate is supported along the entire path rather than only at its endpoint. Reporting both directions is informative because forward and backward work probe the same window from opposite ensembles: agreement in both directions indicates that the overlap is genuinely balanced, rather than a one-sided coincidence. Consistency in both phases matters as well, since the final ΔΔ*G* is obtained as the difference between two accumulated free-energy contributions, so an apparently accurate result may arise from error cancellation and thereby mask substantial errors in either phase.

This window-level view is the practical value of interpretability: rather than trusting or rejecting an opaque number, a scientist can see where an estimate comes from and locate the specific window or phase responsible for a questionable result. The W1 distance, expressed in thermal-energy units *RT*, quantifies how far each predicted distribution along the *λ* schedule must be transported to match the corresponding reference FEP distribution, providing a continuous local diagnostic instead of only a global pass/fail verdict. Panels c and d of Fig. 8 make this concrete: panel c enlarges a single window where the four phase–direction curves can be compared directly, while panel d places the same comparison along all 23 windows so that an isolated anomaly is visible against the surrounding path. Such a window can then be escalated to explicit FEP, structural re-analysis, or experiment—precisely the information a scalar predictor cannot provide. The figure also shows the thermodynamic cycle and atom mapping on which the estimate is built.

**Figure 8.**
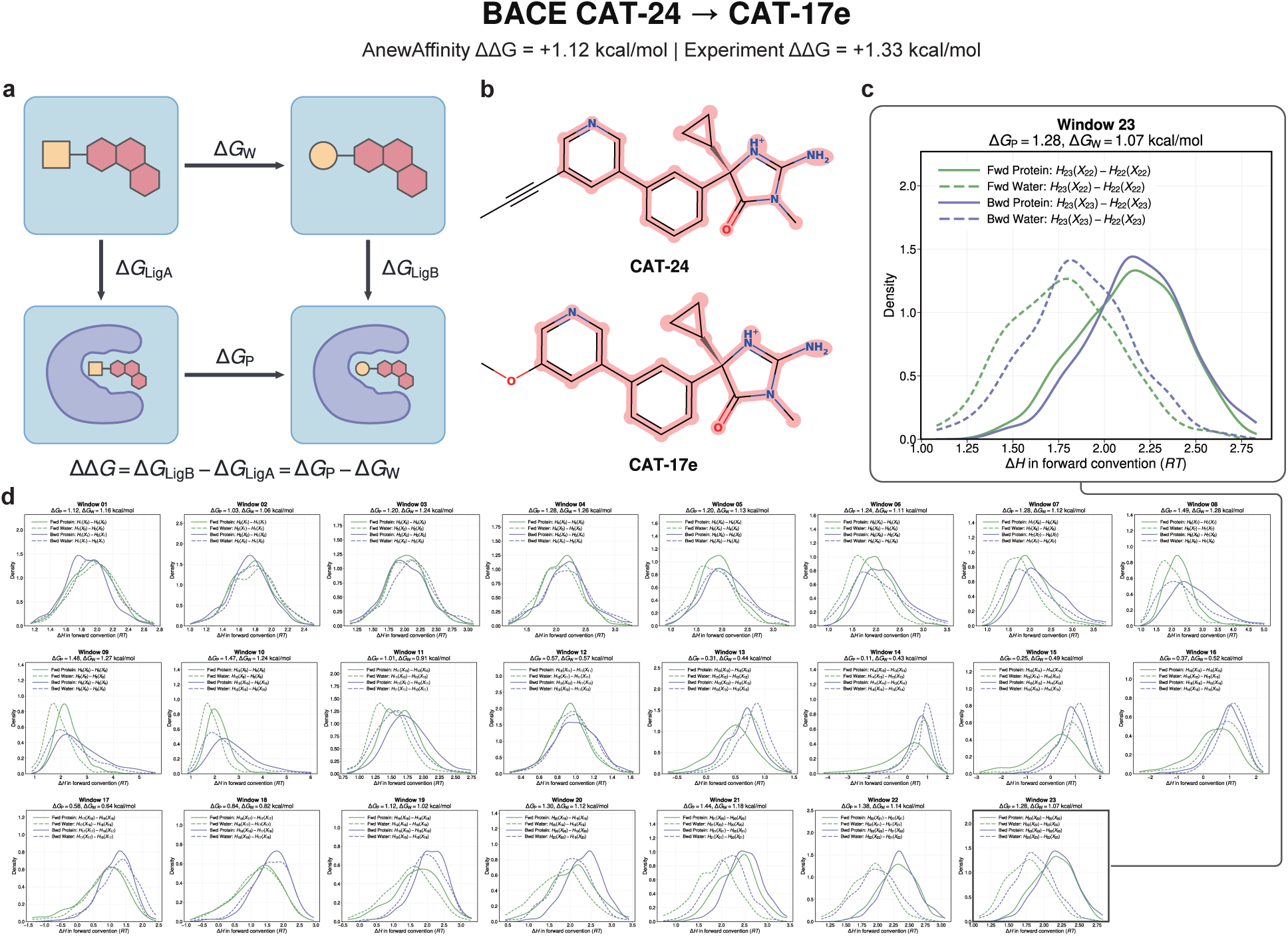
Thermodynamic-cycle estimation for the BACE CAT-24 *→* CAT-17e transformation. **(a)** Protein- and water-phase free energies combine as ΔΔ*G* = Δ*G*_P_ *−* Δ*G*_W_. **(b)** Atom mapping defines the common atoms and changing groups used in both phases. **(c)** Enlarged Window 23 distributions. Green curves show forward work, Δ*H*_fwd_ = *H*_23_(*X*_22_) *− H*_22_(*X*_22_); purple curves show backward samples reflected into the forward convention, *−*Δ*H*_bwd_ = *H*_23_(*X*_23_) *− H*_22_(*X*_23_). Solid and dashed curves denote the protein and water phases, respectively. **(d)** Corresponding distributions across all 23 adjacent *λ* windows, arranged in three rows; dashed guides connect Window 23 to the enlarged view. Values above each window report the protein- and water-phase estimates. BAR is evaluated from the original, unreflected samples, and window estimates are accumulated through the thermodynamic cycle. The resulting AnewAffinity ΔΔ*G* is +1.12 kcal/mol, compared with the experimental value of +1.33 kcal/mol.

Taken together, the benchmark, runtime analysis, and BACE example establish AnewAffinity as a fast, accurate, physically grounded, and interpretable affinity predictor. Its practical distinction is that second-scale estimates are accompanied by phase- and window-resolved evidence, allowing broad candidate sets to be ranked while high-value or ambiguous transformations are identified for escalation to explicit FEP, structural analysis, or experiment. AnewAffinity thereby serves as the quantitative screening layer of AnewDDE, linking affinity prediction directly to the next design round.

## 4 Biologics Design with AnewDesign

Biologics, especially antibodies, stand as one of the most effective modalities of therapeutics in modern medicine. Their widespread adoption across oncology, immunology, infectious disease, and rare genetic disorders is driven by their highly modular architecture, exceptional target specificity, and prolonged systemic half-life [38, 39]. Despite their immense clinical potential, traditional antibody discovery campaigns remain severely bottle-necked by slow, labor-intensive pipelines. Conventional generation relies on camelid immunization and *in vitro* display technologies, which necessitate the construction of massive libraries containing billions of variants, multiple iterative enrichment rounds, and arduous target-specific optimization to yield viable candidates. Recently, a wave of advanced deep learning architectures – spanning all-atom diffusion frameworks, inverse-folding, structural hallucination, and protein language models – has begun to bypass these empirical hurdles by designing high-affinity epitope-specific antibodies or nanobodies [8, 40–54]. Despite rapid advances in generative protein design, a critical bottleneck persists: most computationally designed binders initially exhibit only modest binding affinities in the range from tens to hundreds of nanomolars, ultimately falling back on extensive experimental screening or iterative *in vitro* affinity maturation to achieve therapeutic relevance. Generating true high-affinity (e.g., single-digit nanomolar *K_D_*) binding is fundamentally more challenging than simply predicting plausible structural interfaces.

Building upon the above-mentioned generative paradigm, we introduce AnewDesign, an agentic lab-in-the-loop workflow for the *de novo* generation and *in silico* optimization of therapeutic antibodies and nanobodies, as shown in Fig. 9. It runs on frontier language models in a web browser, so a scientist can easily steer a design campaign in natural language. In this report, we demonstrate a real-world nanobody design campaign using AnewDesign, including *de novo* nanobody design, wet-lab data analysis, as well as *in silico* affinity maturation to reach single-digit nanomolar *K_D_* as measured by multi-concentration SPR on a Carterra LSA XT instrument. This streamlined approach fundamentally collapses conventional discovery timelines, reducing the requisite experimental burden from billions of random variants to a concise panel (i.e., 100-200 clones) of high-confidence and ready-to-test candidates.

**Figure 9.**
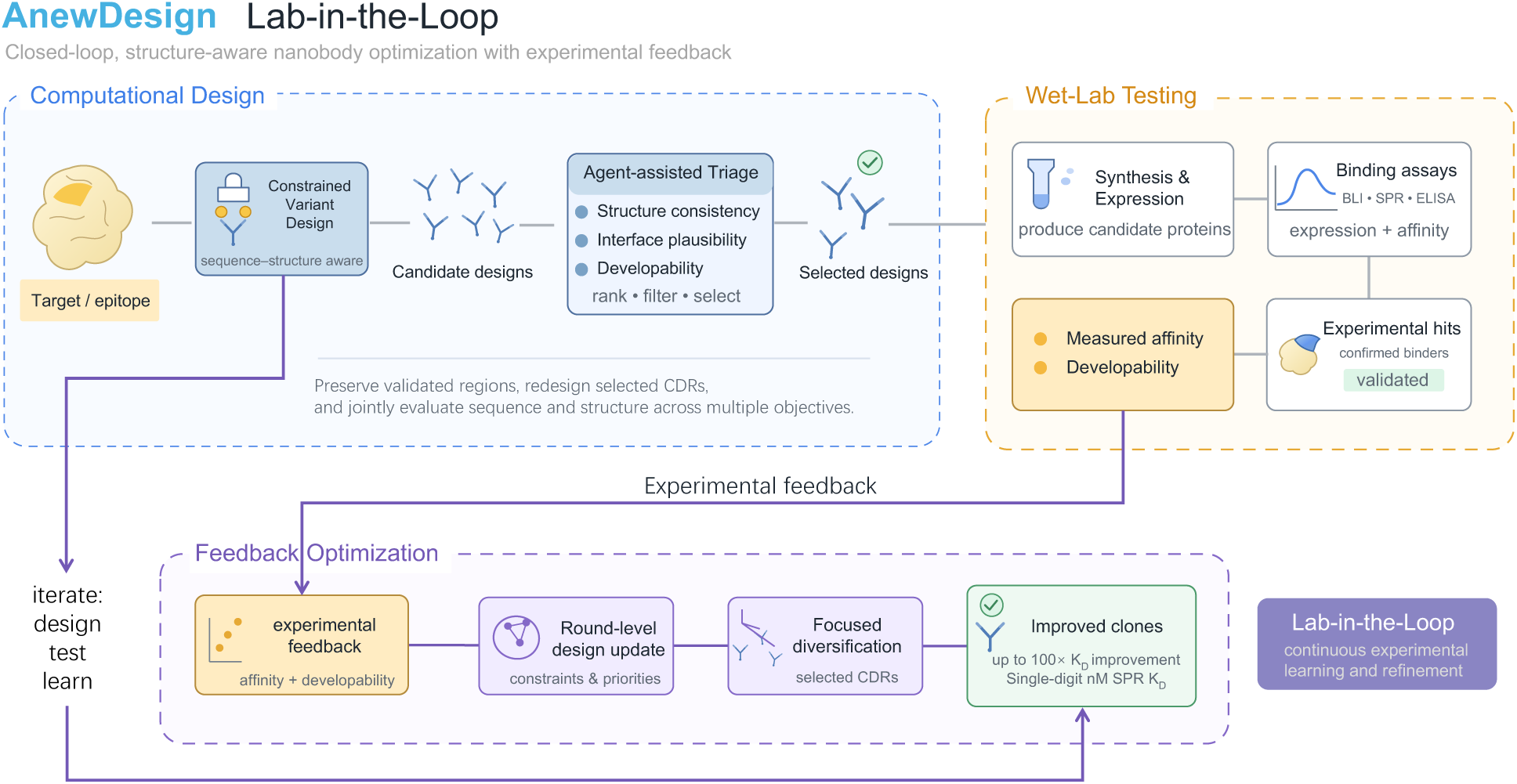
The AnewDesign Lab-in-the-Loop Workflow. This closed-loop, structure-aware platform integrates computational generation with continuous experimental feedback to systematically optimize nanobody candidates. The iterative “design-test-learn” cycle consists of three core phases. *Computational Design*: Starting from a defined target with a set of epitopes, a deep-learning-based algorithm generates a pool of initial candidates with preferred protein-protein interactions. An agent-assisted triage process then ranks, filters, and selects designs based on structural consistency, interface plausibility, and developability. *Wet-Lab Testing*: The selected computational designs are expressed and purified. Candidate proteins undergo wet-lab validation through binding (e.g., BLI, SPR, ELISA) and developability assays. *Feedback Optimization*: Experimental results are fed directly back into the agentic workflow to update round-level design constraints and priorities. By preserving validated regions and applying a focused *in silico* affinity maturation algorithm to selected CDR and nearby framework residues, the workflow generates iteratively improved clones with better binding affinity. This continuous learning cycle can quickly drive up to a 100-fold improvement in binding affinity, achieving single-digit nanomolar *K_D_* values.

To demonstrate the AnewDesign workflow, we conducted a proof-of-concept nanobody design campaign on an internal protein target, as illustrated in Fig. 10. Given the objective of designing VHH binders, the agent begins by constructing the context relevant to biologics design. It first recommends initiating from experimentally reported VHHs to assess their binding epitopes. Guided by this rationale, the agent then retrieves relevant literature and patents, queries structural and sequence databases to characterize the target and its biology, identifies structures suitable for design, and proposes candidate epitopes. Epitope selection is filtered against functional criteria and informed by spatial reasoning over the structure, ensuring that the chosen surfaces reflect the intended mechanism rather than merely geometric convenience. In this case, the agent selects a published crystal structure to identify potential epitope residues suitable for VHH design. It also examines a separate structure of the target protein in complex with its binding partner to delineate the protein-protein interface to be blocked, and overlays the candidate epitope onto this interface to confirm its relevance for inhibition. Once the target and epitope are established, the agent initiates targeted computational design experiments, orchestrating the runs required to generate candidate binders against the selected site. The underlying antibody design methodology is computationally driven, combining diffusion-based generative models with discriminative models built on structure prediction.

**Figure 10.**
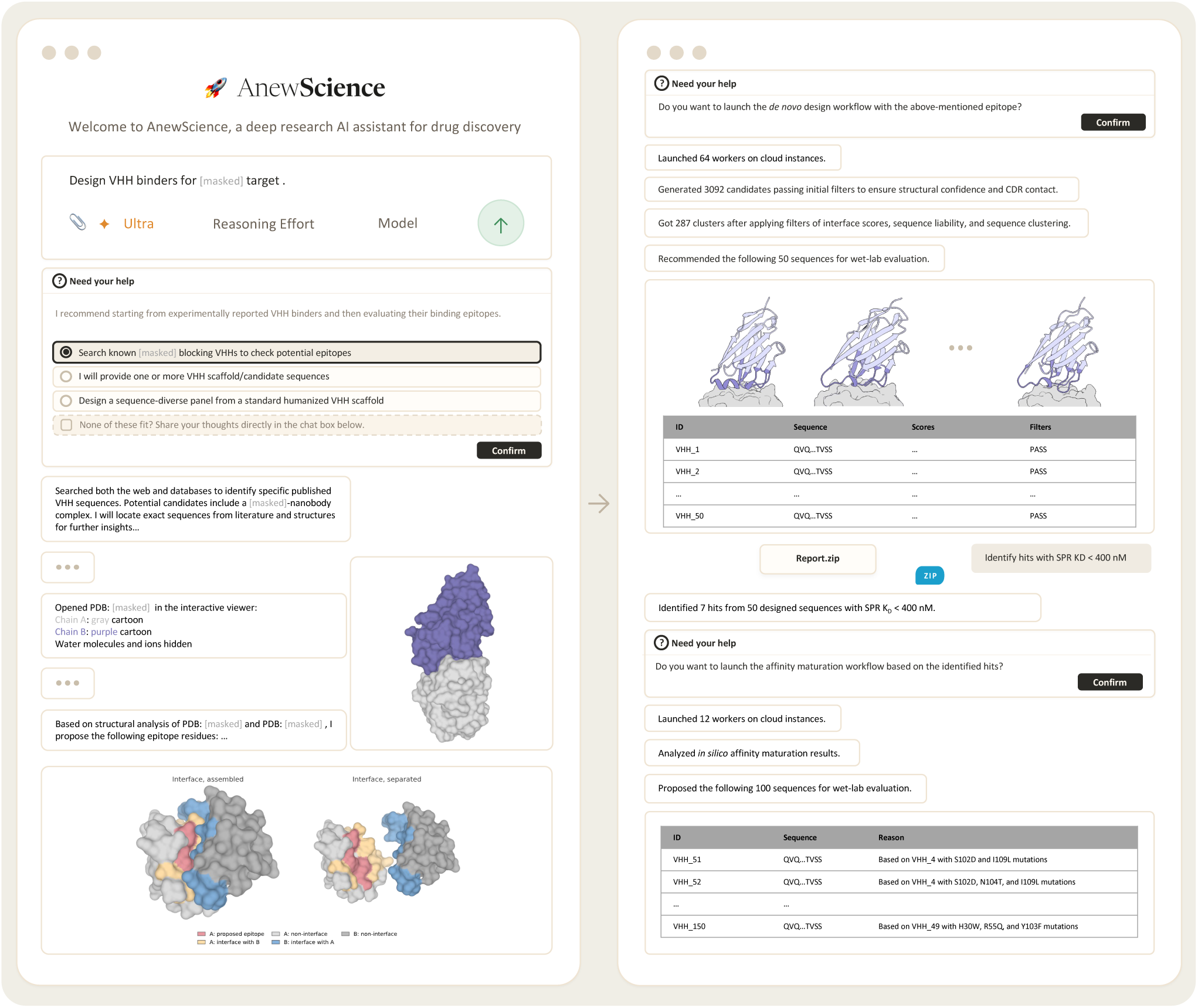
A nanobody design campaign orchestrated by the AnewDesign agentic workflow through AnewScience interface. The left panel illustrates the initial setup where the AI assistant interprets the user’s prompt, autonomously queries structural databases and literature to identify epitopes suitable for the design goal, and prepares the computational design parameters. The right panel demonstrates the execution of the generative pipeline on distributed cloud infrastructure, progressively filtering over 3,000 generated candidates down to a curated panel of 50 *de novo* sequences for the 1st round of wet-lab evaluation. Following the integration of experimental SPR binding data, the assistant automatically identifies hits and then initiates a downstream *in silico* affinity maturation cycle, and finally proposes 100 more candidates for wet-lab evaluation.

The resulting designs are then put through automated quality assurance such as sequence liability and diversity. In particular, we removed sequences with high-risk liabilities such as deamidation, fragmentation, isomerization and N-linked glycosylation by identifying relevant motifs in the CDR regions. The agent also clusters candidates to ensure the set it advances is structurally and sequentially diverse rather than many variations on one solution. What comes out is a curated, lab-ready set rather than a raw pool of model outputs. At the end of the design step, the agent selected 50 VHH sequences for wet-lab validation. Candidates were expressed as VHH-Fc fusions, purified, and evaluated for binding to the target using high-throughput, multi-concentration surface plasmon resonance (SPR) on a Carterra LSA XT instrument (see Supplementary Methods, Sections A.3 and A.4). Of these 50 clones, 7 exhibited SPR-derived *K_D_* values below 400 nM, corresponding to a hit rate of 14% (Fig. 11).

**Figure 11.**
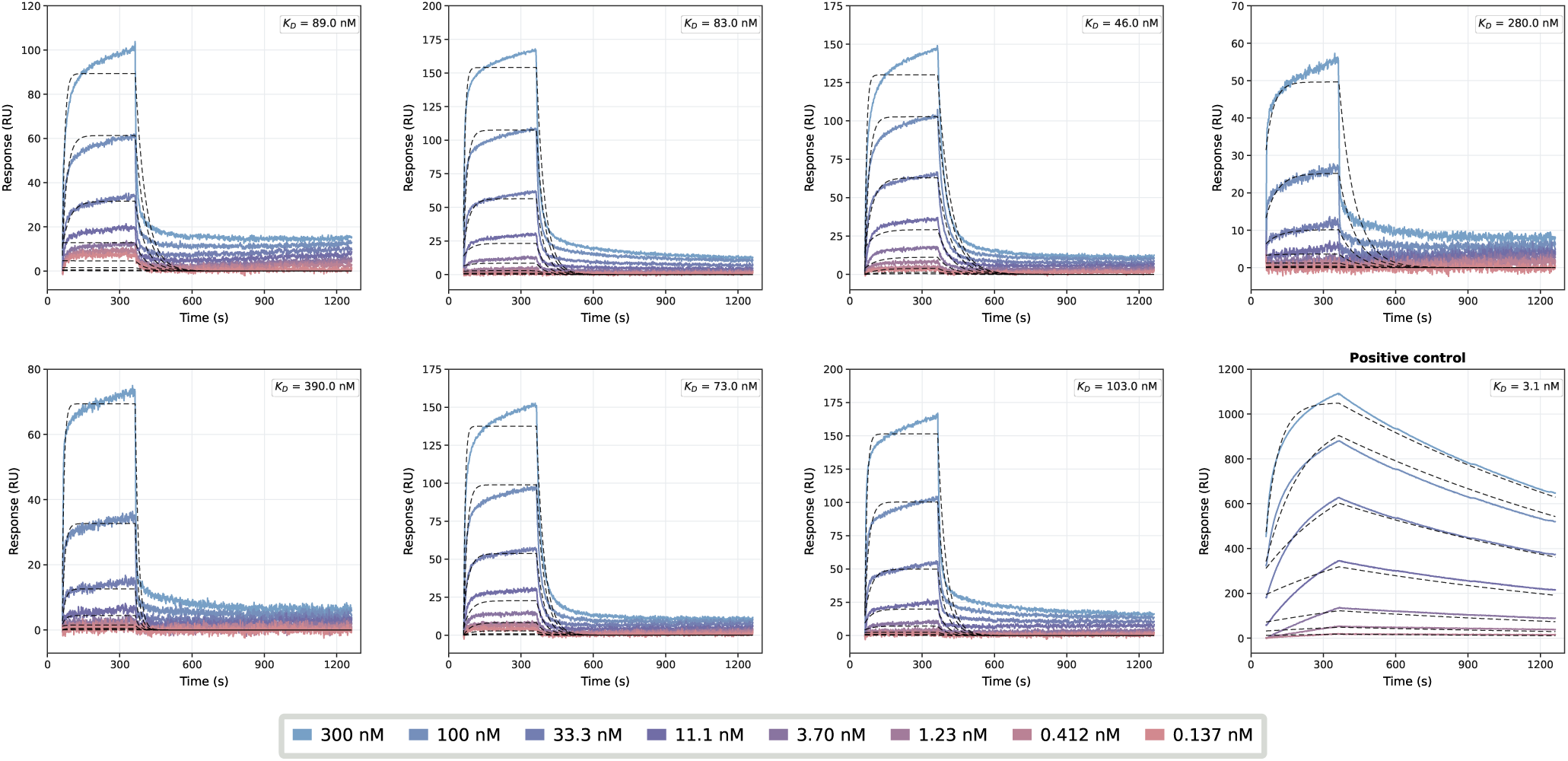
SPR response curves of VHHs *de novo* designed against an internal protein target, with affinities below 400 nM. Binding affinities were measured using up to 8 analyte concentrations (from 300 nM with 3-fold dilution, shown in the bottom colormap) with kinetic fitting to determine *K_D_*. A published nanobody was used as the positive control listed as the last plot. Detailed experimental procedures are provided in Supplementary Methods.

We then move on to the *in silico* affinity maturation stage. The measured affinities and developability labels are used as reference data to further optimize the generative models. Through partial diffusion-based algorithms and an agent-assisted candidate selection pipeline, the selected hits undergo *in silico* affinity maturation to improve both binding affinity and developability before the next round of wet-lab testing. By continuously feeding experimental results back into computational design, we establish a Lab-in-the-Loop framework that enables iterative design, testing, and optimization. The affinity maturation workflow proposed 100 additional clones for experimental validation using the same protein production and SPR assay platform (see Supplementary Methods). This round reduced *K_D_* values by up to two orders of magnitude and yielded 16 clones with single-digit nanomolar *K_D_* values (Fig. 12). These clones are ready for subsequent developability and functional evaluation.

**Figure 12.**
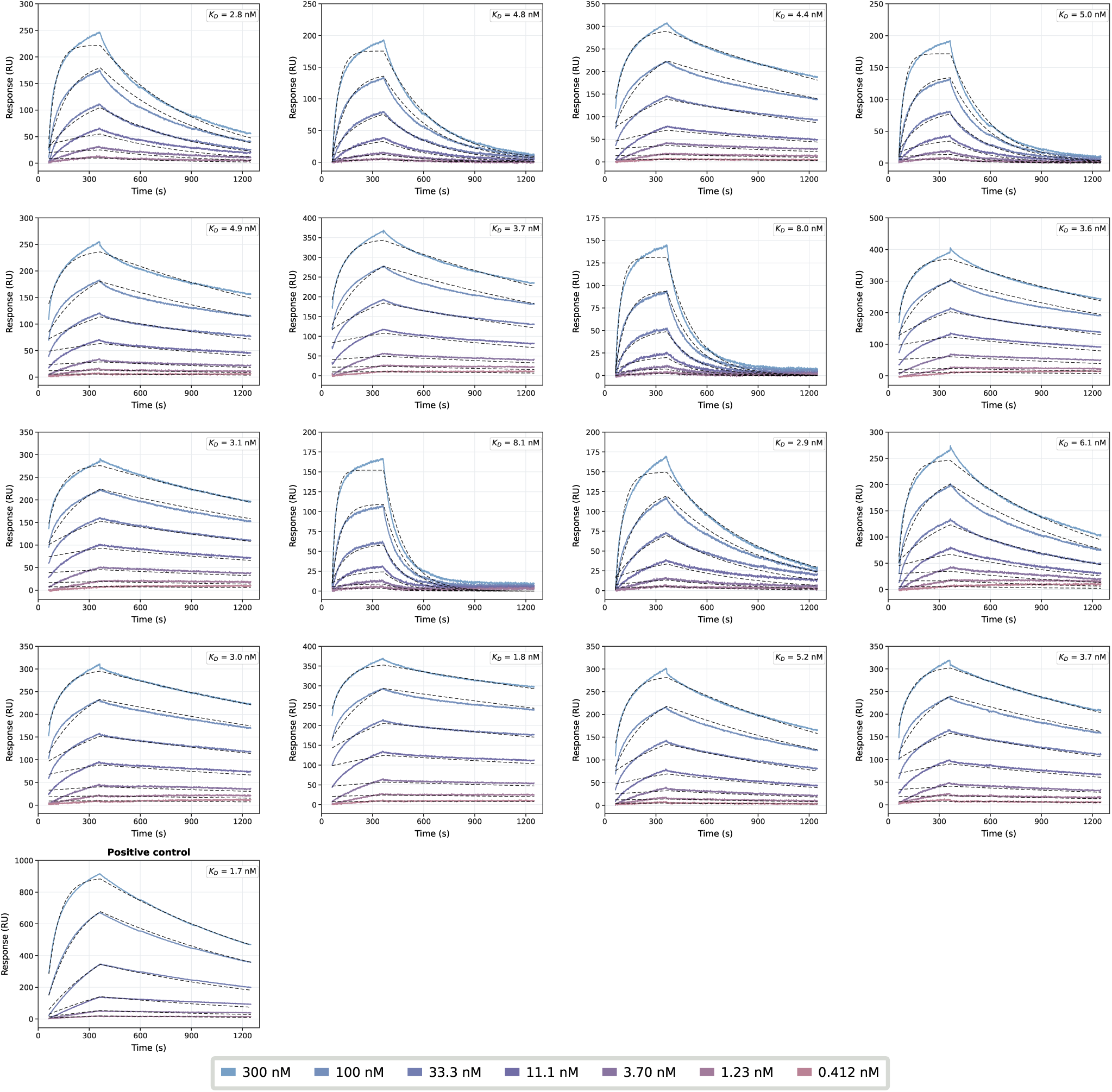
SPR response curves of VHHs against an internal protein target after *in silico* affinity maturation, with affinities below 10 nM. Binding affinities were measured using up to 7 analyte concentrations (from 300 nM with 3-fold dilution, shown in the bottom colormap) with kinetic fitting to determine *K_D_*. A published nanobody was used as the positive control. Detailed experimental procedures are provided in Supplementary Methods.

While this manuscript establishes the foundational architecture of AnewDesign, we present its application to a proof-of-concept internal protein target. This focused demonstration serves as a preview of the platform’s generative capabilities. AnewDesign will be further deployed across a diverse, highly challenging array of disease-relevant targets supported by extensive functional and developability assays to rigorously benchmark the pipeline’s real-world translational potential.

## 5 ADMET and Pharmaceutical Reasoning with AnewMind

Over the past several years, the field of large language models (LLMs) has undergone a profound transformation, driven by major advances in model architectures [55] and large-scale training paradigms [56]. State-of-the-art models now demonstrate unprecedented capabilities in complex open-domain reasoning [57], coding [58], and long-horizon task execution [59]. These advances have not only expanded the frontiers of general-purpose natural language processing, but also provided powerful foundations for scientific research [60] and drug discovery [61].

At Anew, we have regularly evaluated successive generations of state-of-the-art LLMs across a diverse set of drug discovery tasks using our internal benchmarks. These evaluations demonstrate a steady improvement in model capabilities, with particularly rapid progress over the past six months. Because the benchmarks are not publicly available, models are unlikely to have been explicitly optimized for them [62, 63], making the results a useful independent measure of how their underlying capabilities have evolved over time.

Based on this evidence, we believe LLMs are approaching an important turning point: they now have the potential to contribute directly to drug discovery, rather than serving merely as orchestrators that coordinate workflows built around specialized models [64]. Their potential extends beyond analytical tasks such as ADMET prediction [65]. LLMs could also develop into capable R&D scientists with broad domain knowledge and strong independent reasoning abilities, providing practical decision support to pharmaceutical scientists [66].

However, even the most powerful proprietary models may fall short when faced with the practical demands of real-world drug discovery [67]. Meanwhile, modern drug discovery pipelines incorporate a wide range of specialized AI models and computational tools, as well as various wet-lab experiments. We therefore expect the field to evolve toward integrated systems in which the foundation model itself understands end-to-end drug discovery workflows and interacts with real-world experiment feedback [68]. This paradigm enables the continuous generation of high-quality domain-specific data, thereby establishing a feedback loop for further model improvement [69].

Taken together, these considerations make domain-specific post-training of open-source LLM models with strong reasoning capabilities a natural and reasonable approach [70]. Specifically, we conduct full-parameter post-training at the hundred-billion-parameter scale. We report AnewMind Preview’s results on general-purpose benchmarks, ADMET benchmarks, and PharmBench, our internal benchmarks testing capabilities for real world drug discovery R&D pipelines.

### 5.1 General Knowledge Benchmark

The motivation for evaluating general-purpose capabilities is straightforward: a domain-specialized model should enhance scientific and drug-design capabilities without sacrificing general competence [71]. Evaluation on widely adopted general-knowledge benchmarks provides a cost-effective, high-confidence check against domain overfitting. It also verifies that the model retains broad world knowledge and the computational, analytical, and reasoning abilities required to solve complex tasks. AnewMind Preview remains broadly comparable to leading general-purpose models on MMLU-Pro and GPQA Diamond benchmark [72, 73]. As shown in Fig. 13, these results suggest that domain-specific post-training preserves the model’s general knowledge and reasoning capabilities.

**Figure 13.**
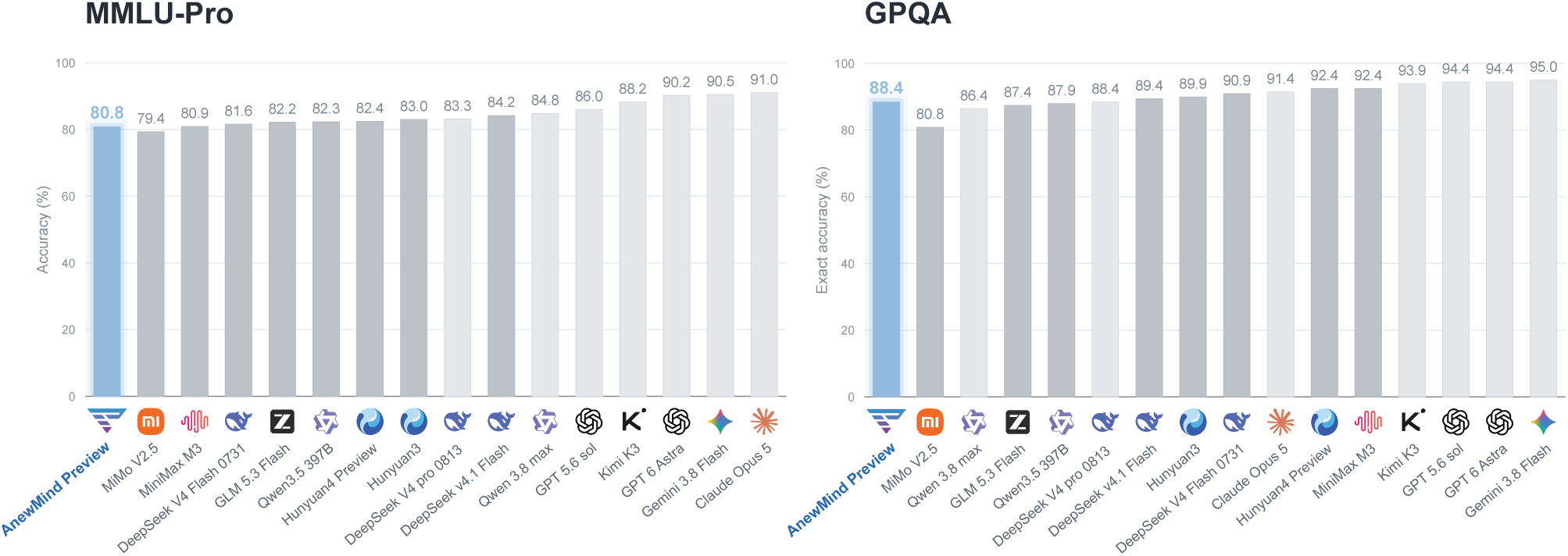
General benchmark performance on MMLU-Pro (left) and GPQA Diamond (right). Bars show accuracy, with AnewMind Preview highlighted in blue and external models shown in gray.

### 5.2 ADMET and Developability Benchmark

ADMET properties are critical considerations in small-molecule drug candidate selection and optimization [74]. For cyclic peptides, membrane permeability is a key developability constraint [75], whereas antibody evaluation emphasizes biophysical properties such as stability, aggregation, and nonspecific binding [76]. Accordingly, we developed a modality-stratified benchmark covering classical ADMET endpoints and modality-specific developability properties across small molecules (Fig. 14a), cyclic peptides (Fig. 14b), and antibodies (Fig. 14c). The detailed composition of each benchmark is described in supplementary section (Table S3, S4).

**Figure 14.**
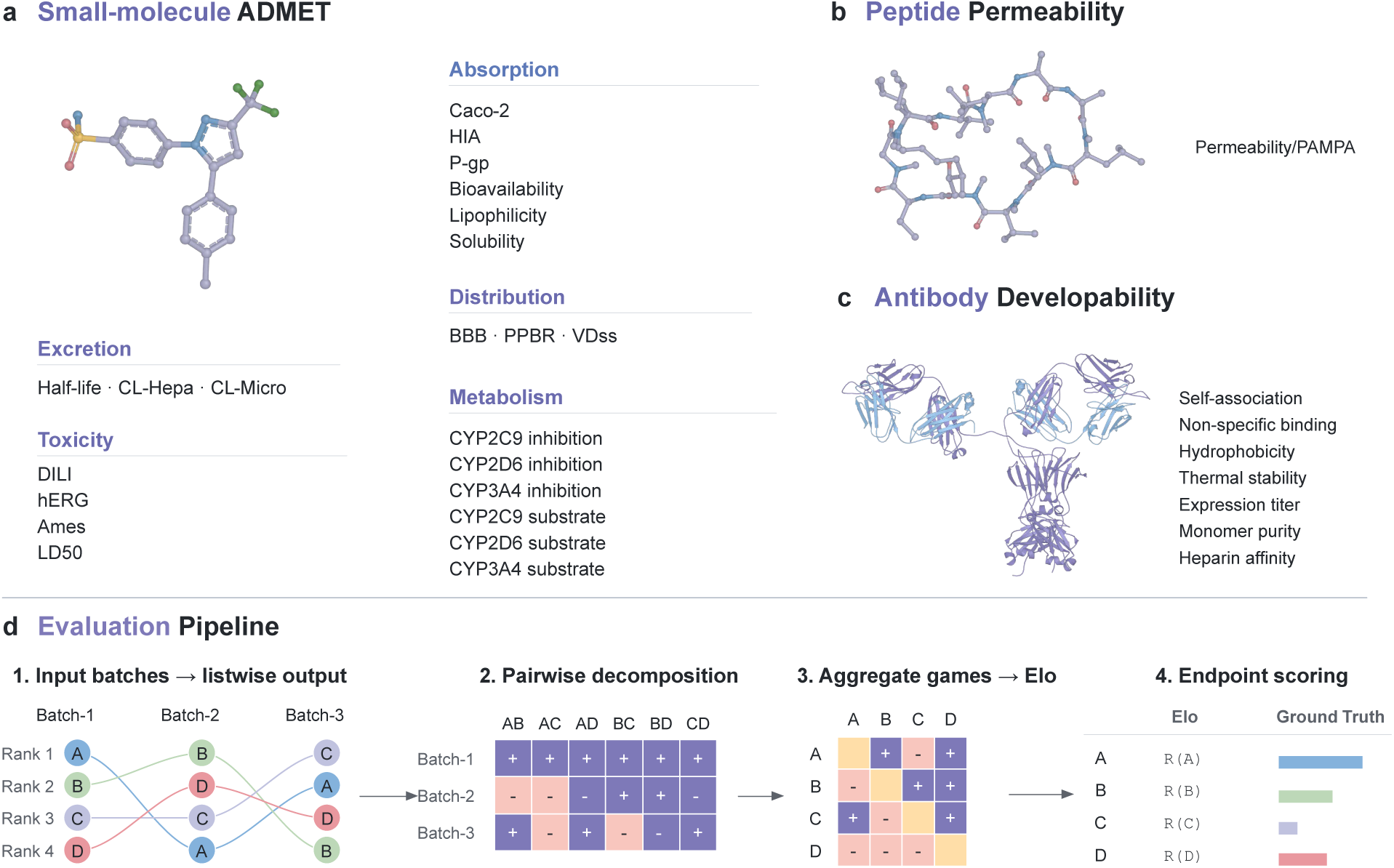
Benchmark scope and evaluation workflow. **(a)** Twenty-two TDC small-molecule endpoints across the five ADMET categories. **(b)** Cyclic-peptide PAMPA permeability. **(c)** Seven antibody developability properties. **(d)** Listwise rankings are decomposed into pairwise outcomes and aggregated into endpoint-specific Elo ratings, which are evaluated against experimental data using AUROC for classification and Spearman’s *ρ* for regression endpoints. Candidate labels A–D are illustrative.

This benchmark does not require LLMs to directly predict absolute experimental measurements. The underlying ADMET datasets span heterogeneous assays, measurement units, numerical scales, and target transformations [77]. Moreover, the general-purpose LLMs evaluated in this study are accessed as text generators rather than trained and calibrated as endpoint-specific regressors. Direct point prediction would therefore confound errors in unit interpretation, scale calibration, and output formatting with errors in capturing structure–property relationships. LLM behavior is also sensitive to prompt formulation and the selection and order of in-context examples [78, 79], as well as to the ordering of candidate options [80]. Under stochastic decoding, repeated inference can produce different reasoning paths or final outputs [81]. Consequently, a single absolute-value prediction may not reliably isolate a model’s ability to identify structure–property trends.

Beyond improving evaluation robustness, relative ranking more closely reflects how predictive models are used in practical drug discovery. In virtual screening, lead optimization, and candidate selection, models are generally used to prioritize candidates under limited synthesis and experimental resources rather than to replace experimental assays with high-precision absolute measurements [82].

Whether optimizing the activity and ADMET profiles of small molecules, screening cyclic peptides for permeability, or assessing antibody developability, the primary decision is which candidates should advance to the next experimental stage. A ranking-based evaluation therefore measures candidate-prioritization performance more directly and better reflects decision-making requirements in real-world drug discovery workflows.

Based on these considerations, each property is formulated as a relative-ranking task (Fig. 14d). Position balancing and repeated scheduling are used to compare each candidate across multiple overlapping candidate sets. The scoring procedure decomposes each listwise ranking into pairwise preference outcomes and aggregates them using the Elo algorithm to obtain a global relative rating within each endpoint [83]. Multiple fixed-seed reorderings of the same pairwise outcomes are performed, and the resulting ratings are averaged to reduce sensitivity to the Elo update order. For classification tasks, ELO-AUROC is calculated using the Elo ratings and experimental class labels [84], for continuous endpoints, Spearman’s rank correlation coefficient is calculated between the Elo ratings and experimental measurements [85]. This framework focuses on evaluating a model’s ability to recover experimentally observed candidate orderings for individual properties and support candidate prioritization, while reducing the influence of absolute-value calibration and run-to-run output variability.

#### 5.2.1 Zero-shot ADMET and Developability Performance across Different Drug Modalities

The overall comparison summarizes the zero-shot performance of AnewMind Preview across the four evaluation settings, as shown in Fig. 15. The model achieved a macro-average Elo-AUROC of 0.771 for small-molecule classification, a macro-average Elo-Spearman correlation of 0.520 for small-molecule regression, an Elo-AUROC of 0.737 for cyclic-peptide PAMPA permeability, and a macro-average Elo-Spearman correlation of 0.247 for antibody developability. Based on point estimates, it ranked fifth of 17, fifth of 16, first of 17, and second of 17 models, respectively. Although scores are not directly comparable across panels because different metrics are used, AnewMind Preview ranked among the top five models in all four settings.

**Figure 15.**
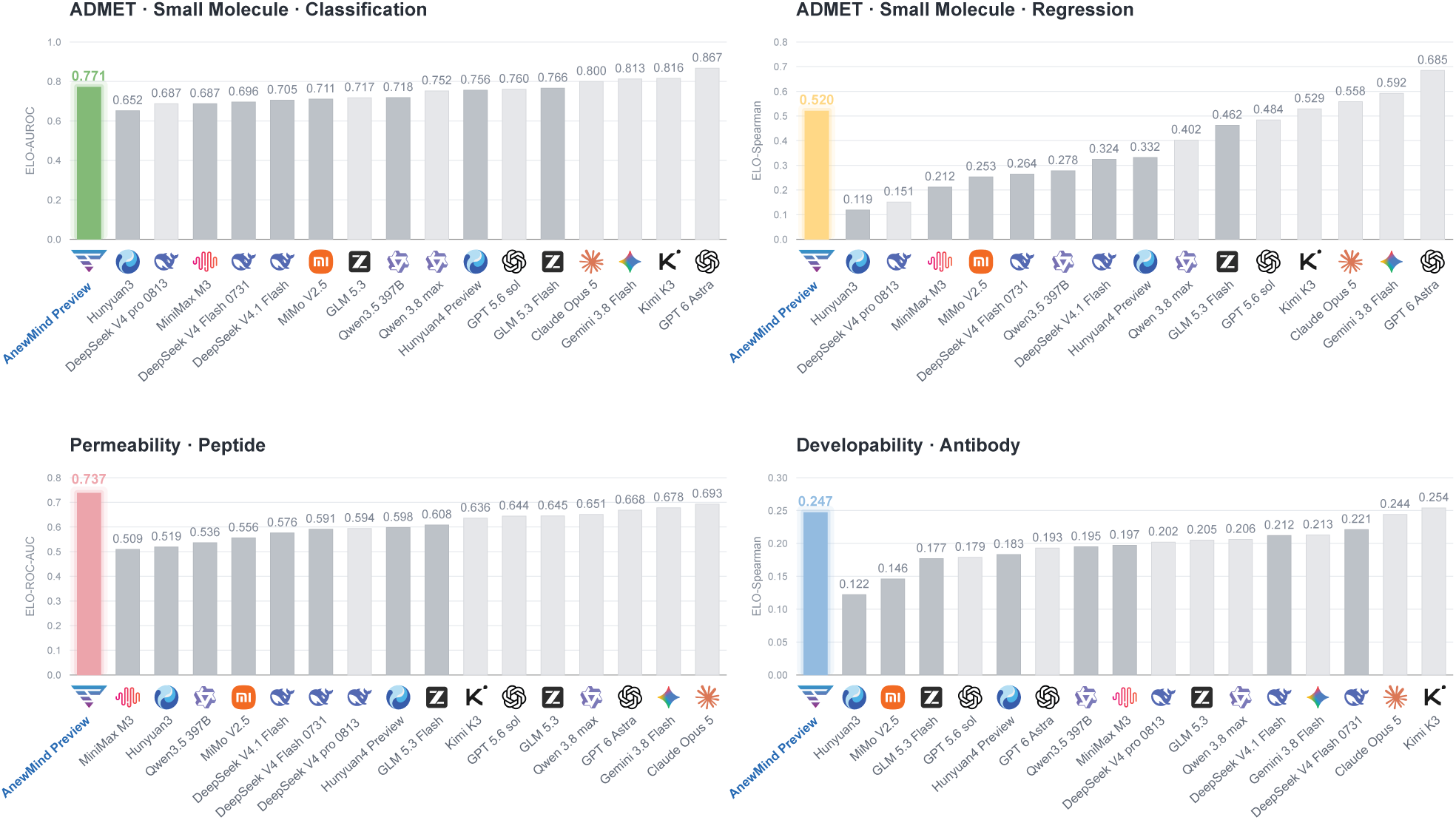
ADMET and developability benchmark. Prediction performance across small-molecule ADMET classification and regression, peptide permeability, and antibody developability. Colored bars highlight AnewMind Preview; higher scores indicate better performance.

Detailed small-molecule results are presented in the supplementary Fig. S2, S3. AnewMind Preview achieved the highest point estimates for CYP2C9, CYP2D6, and CYP3A4 inhibition, with ELO-AUROC values of 0.851, 0.842, and 0.834, exceeding the corresponding second-best results by 0.030, 0.012, and 0.055. Strong classification performance was also observed for human intestinal absorption (0.974), P-glycoprotein inhibition (0.876), blood–brain barrier permeability (0.867), and hERG inhibition (0.862). Among continuous properties, the highest correlations were obtained for aqueous solubility (*ρ* = 0.824), Caco-2 permeability (*ρ* = 0.661), and lipophilicity (*ρ* = 0.607); plasma protein binding (*ρ* = 0.492) was close to the best result of 0.494. Bioavailability, CYP2C9 substrate classification, drug-induced liver injury, and half-life showed comparatively weaker performance and represent priorities for further improvement.

For cyclic-peptide PAMPA permeability, AnewMind Preview achieved the highest Elo-AUROC among the evaluated models (0.737), exceeding the strongest comparator (0.693) by 0.044. This result indicates a clear advantage in prioritizing cyclic peptides with higher passive membrane permeability, as measured by PAMPA.

For antibody developability, AnewMind Preview achieved a macro-averaged Elo-Spearman correlation of 0.247, ranking second among 17 models and trailing the highest-performing model by only 0.007. Its strongest property-level ranking performance was observed for heparin affinity (*ρ* = 0.471), expression titer (*ρ* = 0.372), and hydrophobicity (*ρ* = 0.328). Notably, the model achieved the second-highest point estimates for both non-specific binding and expression titer. Monomer purity was the only property with a negative correlation (*ρ* = *−*0.050), indicating the clearest opportunity for further improvement. Complete property-level results are provided in Supplementary Fig. S4.

Overall, the principal strength of AnewMind Preview lies in its consistency across molecular modalities, rather than being confined to a single molecular class or a limited subset of properties. The model ranked among the top five in all four aggregate evaluations, ranked first in cyclic-peptide permeability, achieved near-leading performance in antibody developability, and obtained the highest point estimates for all three CYP inhibition endpoints. Its performance in human intestinal absorption, membrane transport, hERG inhibition, aqueous solubility, and cyclic-peptide permeability further indicates that the model can identify experimentally meaningful property trends across structurally distinct molecular classes, supporting candidate prioritization in small-molecule ADMET optimization, cyclic-peptide permeability screening, and antibody developability assessment.

#### 5.2.2 ADMET Reasoning Case Studies from a Recent Patent

To minimize the risk of overlap between the evaluation data and model-training corpora, we curated a small-molecule ADMET evaluation set from the recently published international patent WO 2026/002036 A1 [86]. The data were extracted from the pharmacokinetic and in vitro liver-microsomal experiments reported in the patent. Published on 2 January 2026, the patent provides a recent data source for evaluating the ability of models to analyze newly disclosed medicinal-chemistry evidence.

Four matched-pair cases were selected: mouse microsomal intrinsic clearance (Fig. 16), human microsomal intrinsic clearance (Fig. S5), mouse plasma Cmax (Fig. S6), and mouse plasma half-life (Fig. S7). Together, these endpoints assess in vitro metabolic stability across two species, peak systemic exposure following oral administration, and in vivo elimination behavior. The models received the candidate molecular structures, assay context, and endpoint-specific preference direction, but not the experimental measurements. They were required to independently rank the candidates and provide a medicinal-chemistry rationale.

**Figure 16.**
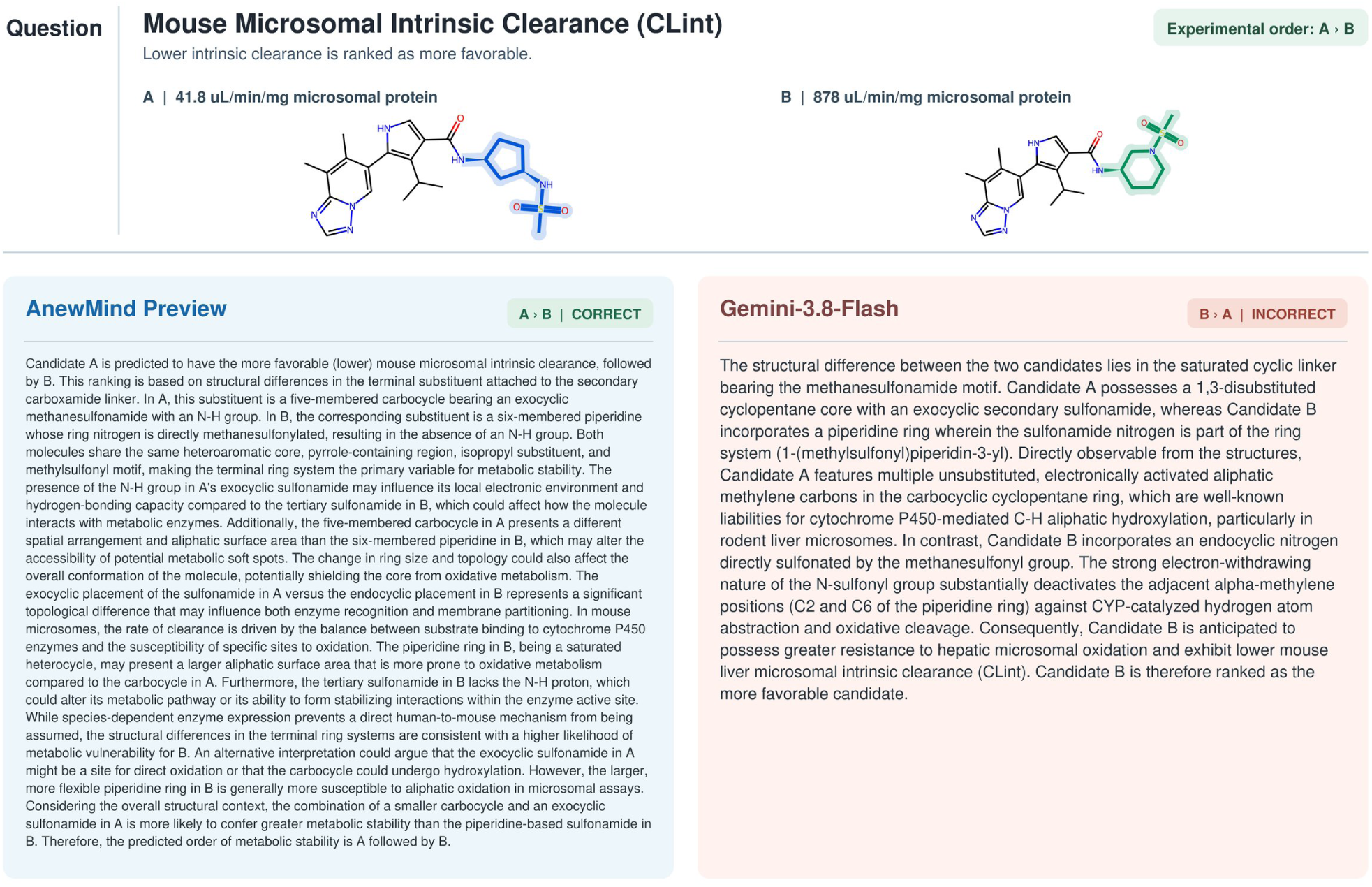
Mouse microsomal intrinsic clearance case study. AnewMind Preview correctly identified the experimental ranking and provided a structure-grounded ADMET rationale, whereas Gemini-3.8-Flash reversed the ranking.

AnewMind Preview reproduced the experimental ranking in all four cases, achieving an accuracy of (4/4). It correctly selected the compounds with lower human and mouse microsomal intrinsic clearance, the compound with higher mouse plasma Cmax, and the compound with a longer mouse plasma half-life. Among the four displayed external-model responses, only GPT-6-Astra correctly ranked the mouse plasma half-life pair; Gemini-3.8-Flash reversed both microsomal-clearance rankings, while GPT-5.6-Sol reversed the Cmax ranking. Manual review further showed that AnewMind Preview accurately identified the principal differences in ring systems, amide environments, fluorinated side chains, and sulfonamide substitution patterns. It integrated these structural observations with appropriately qualified discussions of metabolic stability, absorption, distribution, and clearance. And no significant errors in the identification of structures or functional groups were observed. These examples demonstrate that the model can not only rank matched molecular pairs but also provide structure-grounded explanations that may support compound-progression decisions and the planning of subsequent optimization steps.

### 5.3 PharmBench and Scientific Decision Support

Specialized tools for structure prediction, molecular generation, and ADMET prediction can support well-defined tasks [5, 87, 88]. The broader challenge in drug R&D, however, lies in expert-level reasoning across the development pipeline: integrating heterogeneous experimental findings, explaining changes in compound properties, and making informed trade-offs among competing objectives such as potency, selectivity, pharmacokinetics, and safety. We therefore seek to determine whether a model, once equipped with broad knowledge of drug properties and pharmaceutical R&D, can draw on that knowledge and its reasoning capabilities to analyze real-world problems, propose evidence-based and testable optimization strategies, and design context-specific experimental plans. Moreover, because drug-pipeline information is highly sensitive, models that can be deployed privately and remain under full organizational control are an important requirement for this application.

To evaluate these capabilities, we developed PharmBench, a benchmark designed around authentic drug R&D scenarios (Fig. 17). Our in-house experts authored each question by systematically integrating relevant assays, experimental results, and the conclusions supported by those results, while also incorporating the progression of the R&D scenario and its key decision points. The resulting questions organize otherwise fragmented pieces of evidence within explicit R&D contexts and around specific decision objectives, with the aim of reproducing the judgments and trade-offs that researchers encounter in practice.

**Figure 17.**
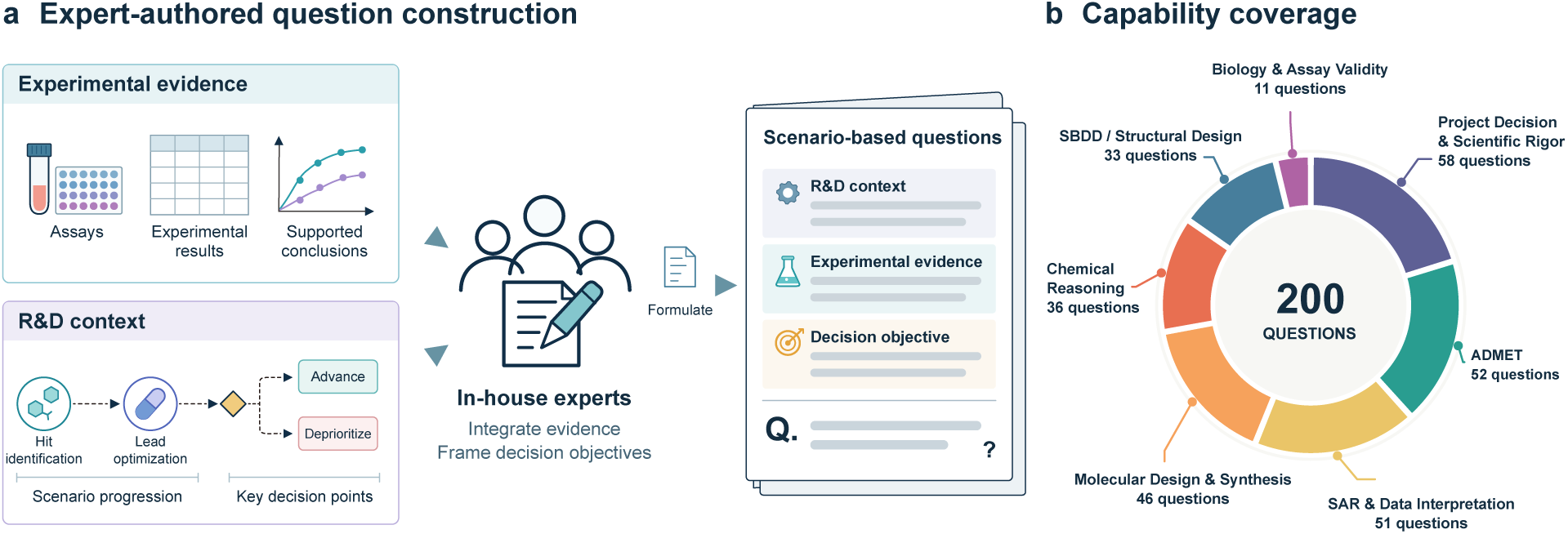
Construction and capability coverage of PharmBench. **(a)** Experts integrate experimental evidence and R&D context to formulate scenario-based questions. **(b)** The 200 subquestions span seven capability dimensions, comprising 287 capability assignments because individual subquestions may carry multiple labels.

PharmBench comprises 50 R&D scenarios (including 200 questions), spanning seven interrelated capability domains. Project decision-making and scientific rigor constitute the most widely represented domain, assessed in 29.0% of the questions, followed by ADMET analysis (26.0%), SAR and data interpretation (25.5%), molecular design and synthesis (23.0%), chemical reasoning (18.0%), structure-based drug design and structural analysis (16.5%), and biology and assay validity (5.5%). Because individual questions may assess multiple capabilities, these percentages are non-exclusive and therefore do not sum to 100%. This design allows the evaluation to focus more directly on a model’s ability to interpret, integrate, and reason over the evidence provided.

When constructing PharmBench, we based the rubrics exclusively on experiments that had been conducted and their observed outcomes. Although some questions may admit other reasonable solutions, this design ensures an objective, traceable, and verifiable basis for evaluation. To further assess a model’s ability to work through continuous project scenarios, we adopted a progressive-disclosure strategy: the model first receives the project background and the initial question, with each subsequent question revealed only after the preceding response is completed. This process simulates the stepwise analysis and decision-making driven by emerging evidence in real-world R&D projects, while preventing results or conclusions from later questions from leaking answers to earlier ones.

In the PharmBench evaluation, AnewMind Preview achieved an overall score of 54.65%, ranking third among 17 models, behind top-ranked GPT-5.6-Sol and only 0.23 percentage points behind second-ranked Kimi K3 (Fig. 18). It exceeded the average score of the 16 external models by 6.59 percentage points and the average score of the eight external hundred-billion-scale models by 10.15 percentage points. AnewMind Preview’s strongest advantage was observed in Molecular Design & Synthesis. Across 46 relevant subquestions, it achieved a score of 60.83%, ranking first among all 17 models, slightly ahead of GPT-6-Astra (60.64%) and GPT-5.6-Sol (60.40%). In ADMET, AnewMind Preview scored 53.11%, ranking fourth and exceeding the external-model average by 7.16 percentage points. Overall, it scored above the external-model average in five of the seven capability dimensions.

**Figure 18.**
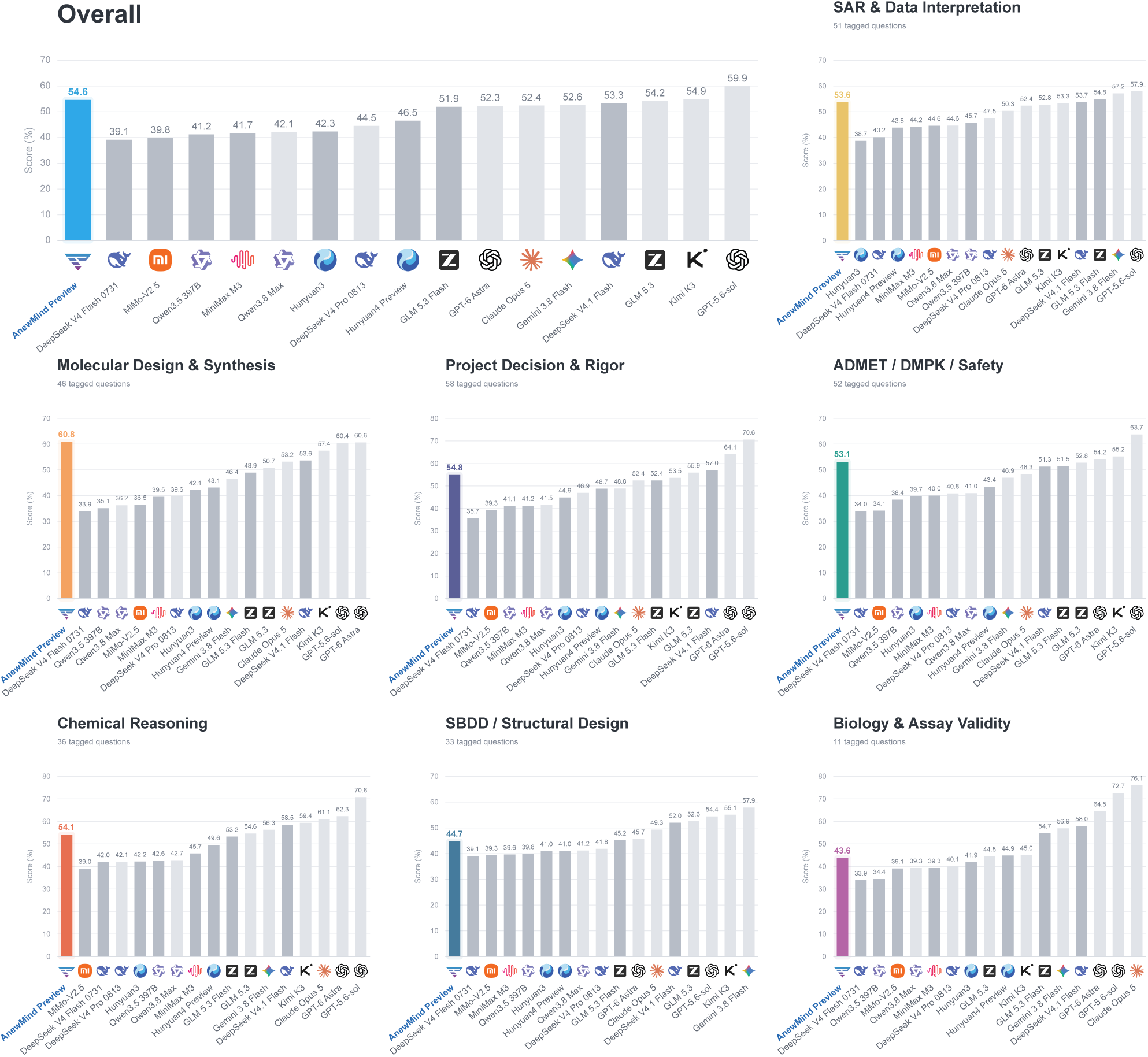
Overall and capability-level performance on PharmBench. Bar charts show weighted scores across 17 models, with AnewMind Preview highlighted in color.

The case-level results further show that AnewMind Preview exceeded the external-model average in 29 of the 50 cases (58%) and achieved the highest score, either outright or jointly, in 7 cases (14%; Fig. 19).

**Figure 19.**
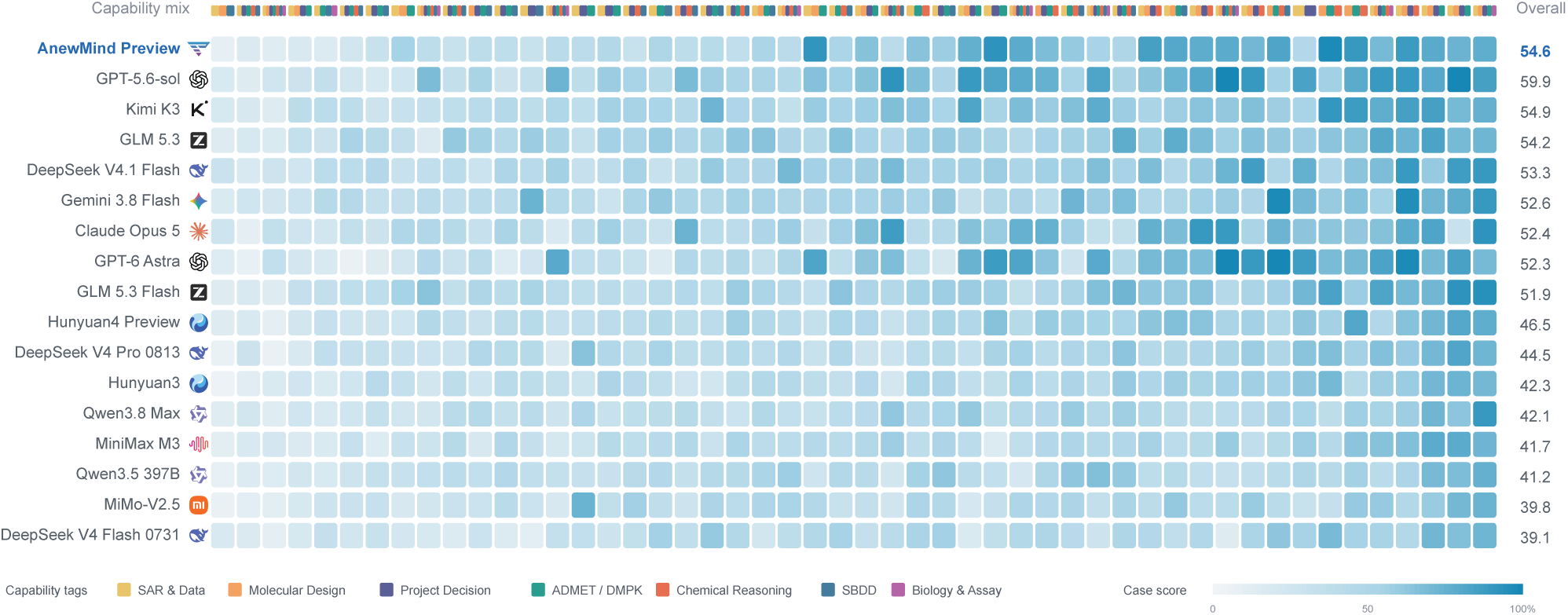
Case-level performance on PharmBench. The heatmap shows 50 cases ordered from hardest to easiest by the mean model score; darker cells indicate higher scores.

To illustrate the task format of PharmBench and the performance of evaluated models, we present representative case studies from small-molecule and antibody development^1^, including the problem context, key experimental evidence, and model responses (Fig. 20). We focus in particular on how models analyze evidence and formulate recommendations for subsequent experiments.

**Figure 20.**
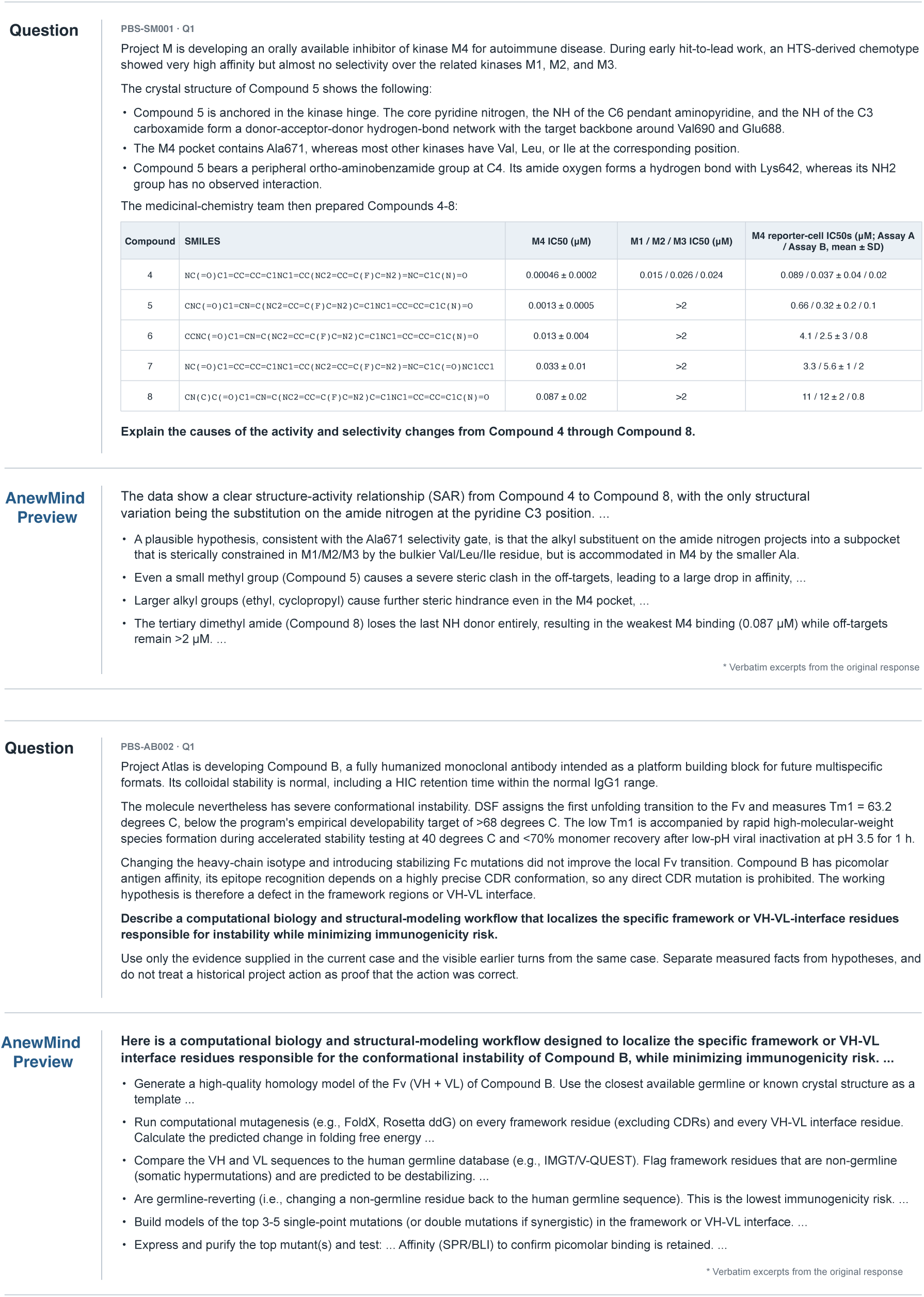
PharmBench case study on kinase inhibitor potency and selectivity (upper) and antibody developability (lower). Ellipses indicate omitted text.

In the small-molecule example, AnewMind Preview interpreted compound SMILES, key crystal-structure interactions, pocket residues, and kinase activity and selectivity data. It derived a coherent structure–activity relationship: C3 amide substitution exploits the smaller ALA671 in M4 pocket, whereas the bulkier VAL, LEU, or ILE residues in related kinases create steric exclusion. The improved selectivity therefore results primarily from weaker off-target binding rather than stronger M4 binding. Further enlargement of the substituent eventually compromises its fit within M4, while removal of the amide NH disrupts a key hinge-binding hydrogen-bond network. By connecting structural differences with activity trends, the model produced a mechanistically grounded SAR hypothesis that medicinal chemists could review and use to inform subsequent optimization.

The antibody example presents a different development challenge. Although the antibody has normal colloidal stability and picomolar antigen affinity, its first Fv unfolding transition occurs at only 63.2°C, below the project target of 68°C, and is accompanied by accelerated aggregation and poor monomer recovery after low-pH treatment. Given the working hypothesis stated in the prompt, AnewMind Preview proposed a workflow to prioritize framework and VH-VL-interface mutations for experimental testing. It proposed combining human-germline residue conservation with three-dimensional analyses of local energetics, packing, steric clashes, and interface quality to prioritize a small set of experimentally testable mutations. Importantly, it treated sequence frequency and computational energy estimates as filters rather than substitutes for experimental validation.

Together, these examples show that AnewMind Preview can support the full reasoning chain from evidence integration and mechanistic hypothesis generation to design recommendations and experimental validation. Its natural-language outputs make assumptions, trade-offs, and uncertainties explicit, allowing medicinal chemists and antibody engineers to efficiently review and refine the analysis. Rather than replacing expert judgment, the model can help researchers organize complex project information, identify overlooked explanations, compare design options, and formulate testable next steps. Combined with its quantitative performance, these results demonstrate AnewMind Preview’s ability to convert complex scientific evidence into interpretable, reviewable, and actionable reasoning under realistic R&D constraints.

## A Supplementary Information

**Table S1.** Top-1 success rates (%) across antibody-antigen, protein-ligand, and protein-protein structure-prediction benchmarks. Bold values indicate the best result for each benchmark. AnewFold results and values marked with *∗* were computed in this study; all other values were taken from publicly reported benchmark results.

| Model | Antibody-Antigen |  |  | Protein-Ligand |
| --- | --- | --- | --- | --- |
|  | FoldBench | PXMeter | AF3-AB | FoldBench |
| AF3 | 48.8 | – | 47.0 | 64.9 |
| Boltz-1 | 34.4 | 19.6 | 14.9 | 55.0 |
| OF3p2 | 34.9 | 27.9 | 30.9 | 44.5 |
| Protenix-v1 | 52.3 | 40.2 | 40.4 | 62.8 |
| Protenix-v2 | 65.0 | 49.7 | 53.5 | 64.0 |
| OpenDDE | 70.0 | 51.0 | 55.6(*) | 60.1 |
| IsoDDE | 75.6 | – | – | 76.0 |
| AnewFold | **76.2** | **60.8** | **62.3** | **77.4** |

### A.1 AnewFold Benchmark Construction

AnewFold’s structural training data comprise only PDB entries with release dates earlier than September 30, 2021, with data preparation following the Protenix-v2 pipeline.

#### A.1.1 Pocket Identification

The pocket identification evaluation set strictly follows IsoDDE’s construction procedure: for systems released after January 01, 2024, all protein–ligand structures with resolution better than 5 Å were collected and filtered by the following criteria: pocket residues are uniformly defined as residues with any atom within 5 Å of any ligand heavy atom; ligands must carry CCD codes and contain 6-50 heavy atoms; common crystallization buffers and common ions are removed; a pocket must contain at least 10 residues; only non-chimera protein chains are retained, i.e., chains mapping to multiple UniProt IDs are excluded; to avoid counting the same pocket multiple times, each ligand’s local pocket is expanded into an alignable protein-chain instance: the contacting chain is aligned to its UniProt reference, pocket residues are represented in UniProt-aligned coordinates, and the ligand centroid is computed in this local frame. Instances sharing the same UniProt reference and having spatially proximal ligand centroids are then grouped into a single pocket cluster by single-linkage clustering with a 5 Å distance threshold, yielding the strict pocket test set. In addition, following the CryptoBench [89] standard, we carve a cryptic-pocket subset from this test set to evaluate hidden-pocket prediction.

#### A.1.2 Molecular glue Structural Prediction

Molecular glue (MG) complexes provide a stringent test of structural generalization because the ligand must be positioned together with two cooperative protein-ligand interfaces, often in geometries that are sparsely represented in training data [22]. We evaluated AnewFold on the MG benchmark and restricted the comparison to 131 common cases produced by AnewFold, Boltz-1, Protenix-v1, and Protenix-v2. Performance was measured using DockQ for the protein-protein interface, ligand RMSD for pose accuracy, and LDDT-PLI for local protein-ligand interaction quality. A prediction was counted as jointly successful only when all three criteria were satisfied simultaneously: DockQ *>* 0.23, ligand RMSD *<* 2.0 Å, and LDDT-PLI *>* 0.8.

**Figure S1.**
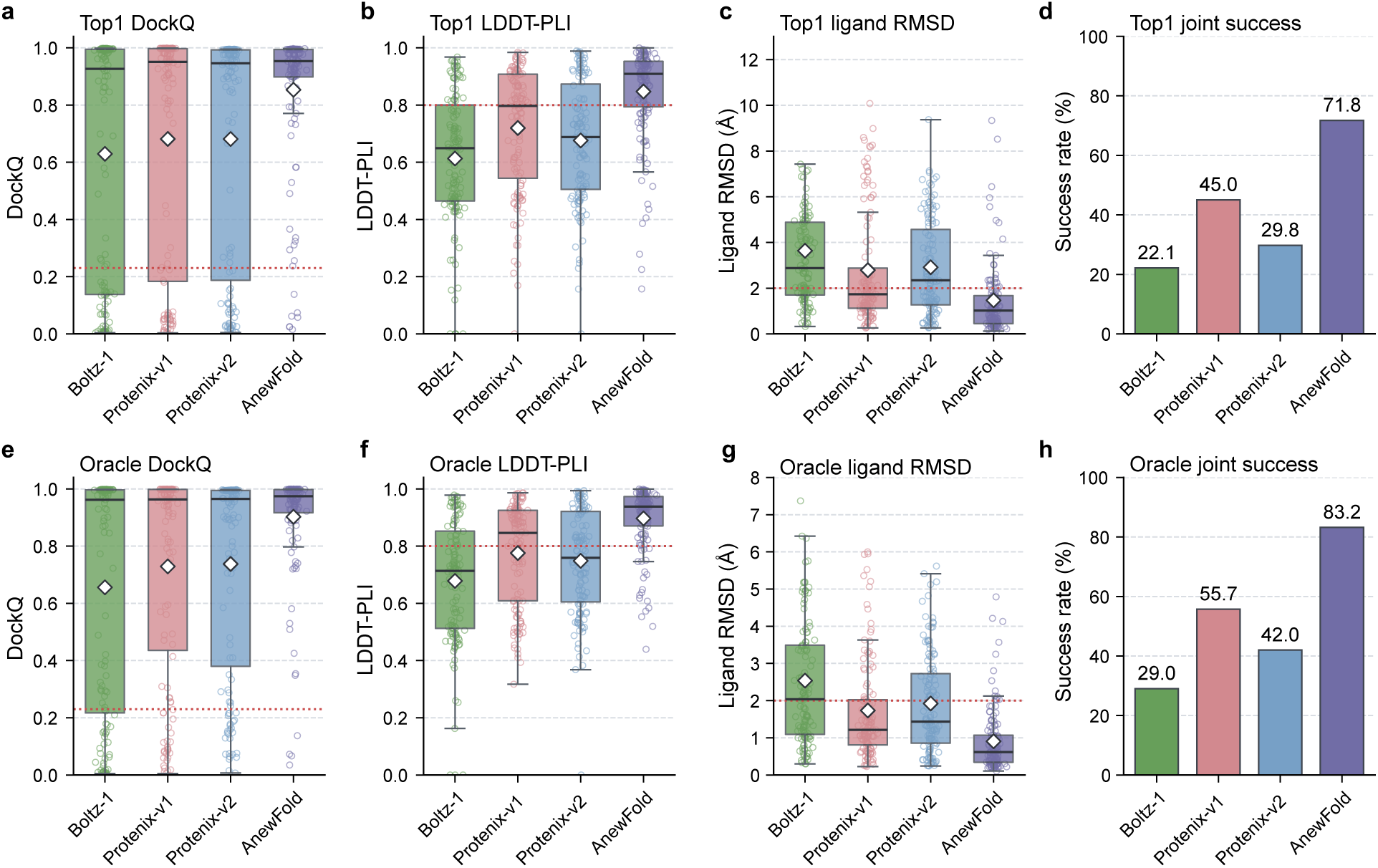
Structural prediction performance on the molecular glue benchmark, evaluated on the 131 complexes for which all four models (AnewFold, Protenix-v1, Protenix-v2, and Boltz-1) produced complete predictions. A prediction counts as a joint success when it satisfies all three criteria simultaneously: DockQ *>* 0.23, ligand RMSD *<* 2.0 Å, and LDDT-PLI *>* 0.8. **(a-d)** Top-1 performance, based on each model’s highest-ranked prediction: distributions of DockQ **(a)**, LDDT-PLI **(b)**, and ligand RMSD **(c)** together with the resulting joint success rate **(d)**. AnewFold achieves a top-1 joint success rate of 71.8%, compared with 45.0% for Protenix-v1, 29.8% for Protenix-v2, and 22.1% for Boltz-1. **(e-h)** Oracle performance over the 25 poses predicted per complex: best DockQ **(e)**, best LDDT-PLI **(f)**, best ligand RMSD **(g)** and oracle joint success **(h)** defined as at least one of the 25 poses satisfying all three criteria. AnewFold reaches an oracle joint success rate of 83.2%, versus 55.7% for Protenix-v1, 42.0% for Protenix-v2, and 29.0% for Boltz-1. Red dashed lines indicate the metric thresholds used in the joint criterion; diamonds denote distribution means. Ligand RMSD is shown with panel-specific upper caps for readability.

Using only the top-ranked prediction, AnewFold attains a 71.8% joint success rate, exceeding Protenix-v1 by 26.8%, Protenix-v2 by 42.0%, and Boltz-1 by 49.7% (Fig. S1a-d). The component-metric distributions are consistent with this result: AnewFold concentrates more predictions above the DockQ and LDDT-PLI thresholds and below the 2.0 Å ligand-RMSD threshold. The gain therefore reflects concurrent improvements in ternary assembly, ligand placement, and local interaction geometry rather than optimization of a single metric.

Sampling 25 poses further improves all methods, but the relative ordering remains unchanged (Fig. S1e-h). AnewFold reaches an oracle joint success rate of 83.2%, compared with 55.7% for Protenix-v1, 42.0% for Protenix-v2, and 29.0% for Boltz-1. The 11.4-percentage-point increase from top-1 to oracle performance indicates that additional sampling recovers difficult conformations, while the already high top-1 rate shows that AnewFold usually ranks a near-native ternary structure first. Together, these results support AnewFold’s ability to model the coupled protein-protein and protein-ligand interactions that stabilize molecular-glue complexes.

### A.2 Affinity Runtime Comparison

**Table S2.** Runtime and Pearson correlation values used in the qualitative runtime-accuracy comparison in Fig. 7. Runtime values span different datasets, hardware, and timing boundaries and therefore do not constitute a controlled speed benchmark. No comparable runtime is reported for IsoDDE, so its value is marked “—” and it is not plotted in Fig. 7; its Pearson correlation of 0.85 is the mean on the FEP+ 4 benchmark reported in the IsoDDE technical report.

| Method | Runtime (s) | Pearson correlation | Method type | Hardware | Value source |
| --- | --- | --- | --- | --- | --- |
| BACPI | 0.00048 | 0.14 | ML | GPU | Boltz-2 study [4] |
| Boltz-2-iptm | 5.0 | 0.04 | ML | GPU | Boltz-2 study [4] |
| Boltz-2 | 20.0 | 0.66 | Boltz-2 | GPU | Boltz-2 study [4] |
| Chemgauss4 (Docking) | 25.0 | 0.26 | Physics | CPU | Estimated from source figure |
| FMO | 300.0 | 0.55 | Physics | CPU | Estimated from source figure |
| MM/PBSA | 720.0 | 0.18 | Physics | CPU | Estimated from source figure |
| OpenFE | 33,000 | 0.66 | Physics | GPU | Gowers et al. [36]; Boltz-2 [4] |
| AnewFEP | 33,000 | 0.77 | Physics | GPU | Estimated from AnewFEP study [34] |
| ABFE | 75,000 | 0.75 | Physics | GPU | Wu et al. [35]; Boltz-2 [4] |
| IsoDDE | — | 0.85 | ML | GPU | IsoDDE technical report [9] |
| AnewAffinity | 3.4 | 0.76 | ML | GPU | Provided measurement |

### A.3 Protein Construction, Expression, and Purification

VHH candidates were formatted as VHH-Fc fusions with a C-terminal human IgG1 Fc (C220S). All recombinant constructs were synthesized and cloned into high-expression mammalian vectors. These proteins were produced using a CHO mammalian cell transient expression system.

Transfected cells were cultured in serum-free media within a humidified incubator (8% CO_2_) at 37*^◦^*C. Between five and seven days following transfection, the culture media was collected, centrifuged, and passed through a 0.22 µm filter to remove cellular debris. To isolate Fc-tagged VHHs, Protein A affinity chromatography was performed utilizing MabSelect resin. The column was washed with PBS (pH 7.4), and the proteins were eluted using a pH 3.5 sodium acetate buffer and promptly neutralized. The recovered proteins were then transferred into either standard PBS or 0.05% PBST via dialysis or buffer exchange. Final protein concentrations were quantified by measuring A280 absorbance on a NanoDrop spectrophotometer. Additionally, sample purity and expected molecular weights were evaluated using CE-SDS, successfully demonstrating the structural integrity of the *∼* 80 kDa VHH-Fc constructs.

### A.4 High-Throughput Multi-Concentration SPR Analysis

High-throughput binding kinetics of the purified nanobodies to target protein were measured using a Carterra LSA XT instrument. The Fc region of a VHH-Fc nanobody is captured or immobilized onto the chip surface (as the ligand) using Protein A/G, and the monomeric antigen is then flowed over it as the mobile phase (analyte). The nanobodies were first immobilized on the sensor chip for 10 minutes. Using 1xHBS EP+ as the running buffer, the antigen was serially diluted from 300 nM with a factor of 3. Antigen injections proceeded in a low-to-high concentration format, preceded by multiple buffer-only injections to establish a stable baseline. Data acquisition for each concentration cycle consisted of a 60-second baseline, 5 minutes for association, and 15 minutes for dissociation. Following each analyte titration series, the chip surface was regenerated using three 20-second pulses of 10 mM Gly-HCl (pH 2.0). Binding kinetics were subsequently calculated using the Carterra KIT™ analysis software applied to a 1:1 Langmuir binding model. A reference channel was included in each cycle for reference subtraction, and the final buffer-only injection immediately preceding the antigen injection was used as the blank for background subtraction of resonance units (RU).

### A.5 ADMET and Developability Benchmark Construction

#### A.5.1 Small Molecule Benchmark

The small-molecule evaluation set follows the official test splits of the TDC ADMET Group [77]. The endpoint composition and dataset size are summarized in the table below.

**Table S3.** Small-molecule evaluation set: ADMET endpoint composition and test-record counts.

| ADMET Category | Classification Endpoints (No. of Test Records) | Regression Endpoints (No. of Test Records) | Subtotal |
| --- | --- | --- | --- |
| Absorption | HIA Hou (117)<br>P-gp Broccatelli (245)<br>Bioavailability Ma (128) | Caco-2 Wang (182)<br>Lipophilicity AstraZeneca (840)<br>Solubility AqSolDB (1,997) | 3,509 |
| Distribution | BBB Martins (406) | PPBR AZ (559)<br>VDss Lombardo (226) | 1,191 |
| Metabolism | CYP2C9/2D6/3A4 Veith inhibition endpoints (2,419/2,626/2,467)<br>CYP2C9/2D6/3A4 Carbon-Mangels substrate endpoints (135 each) | — | 7,917 |
| Excretion | — | Half-life Obach (135)<br>Clearance Microsome AZ (221)<br>Clearance Hepatocyte AZ (243) | 599 |
| Toxicity | hERG (132)<br>Ames (1,457)<br>DILI (96) | LD50 Zhu (1,478) | 3,163 |
| Total | 13 classification endpoints | 9 regression endpoints | 16,379 |

#### A.5.2 Peptide Benchmark

The cyclic-peptide evaluation set was derived from CycPeptMPDB using the PAMPA data curated in Benchmark CycPeptMP [75, 90]. The source data contain 6,311 records, with some cyclic peptides repeated across the original random-split and scaffold-split files. Deduplication based on normalized input hashes resulted in 3,939 unique cyclic peptides. In the current evaluation, the original random and scaffold splits are not treated as separate tasks; instead, the deduplicated samples are combined into a single evaluation set.

#### A.5.3 Antibody Benchmark

The antibody evaluation set was based on GDPa3, the independent test set of the 2025 Ginkgo Datapoints Antibody Developability Competition [91]. GDPa3 contains 80 human IgG1 antibodies derived from the Observed Antibody Space.

Each property was evaluated using all antibodies with a valid experimental measurement, rather than restricting the entire evaluation to the 57 antibodies with complete Core7 data. Of the 80 antibodies, 73 had complete Core5 measurements and 57 had complete Core7 measurements. In total, Core5 and Core7 contained 393 and 534 valid antibody–property measurements, respectively.

**Table S4.**
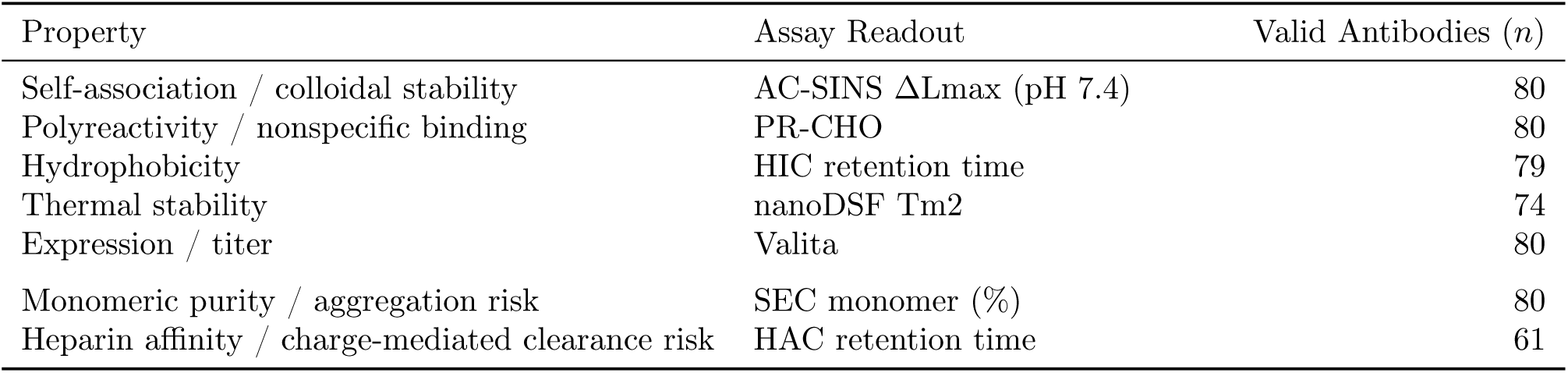
Antibody evaluation set: property composition and valid antibody counts (GDPa3, n = 80).

| Property | Assay Readout | Valid Antibodies ($n$) |
| --- | --- | --- |
| Self-association / colloidal stability | AC-SINS $\Delta L_{\max}$ (pH 7.4) | 80 |
| Polyreactivity / nonspecific binding | PR-CHO | 80 |
| Hydrophobicity | HIC retention time | 79 |
| Thermal stability | nanoDSF $T_{m2}$ | 74 |
| Expression / titer | Valita | 80 |
| Monomeric purity / aggregation risk | SEC monomer (%) | 80 |
| Heparin affinity / charge-mediated clearance risk | HAC retention time | 61 |

### A.6 Performance of Small Molecule ADMET and Antibody Developability Prediction

**Figure S2.**
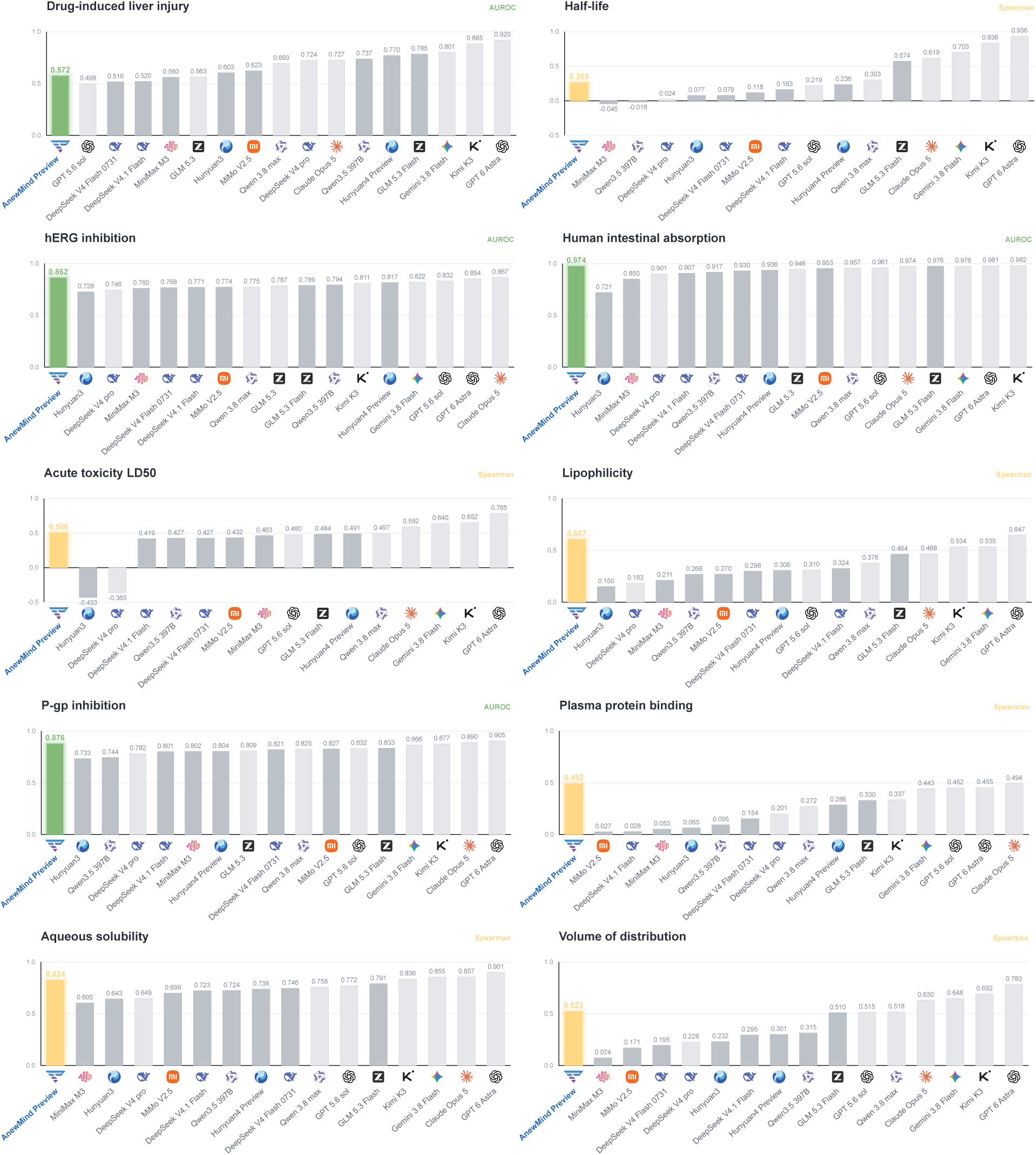
Performance of small molecule ADMET prediction I.

**Figure S3.**
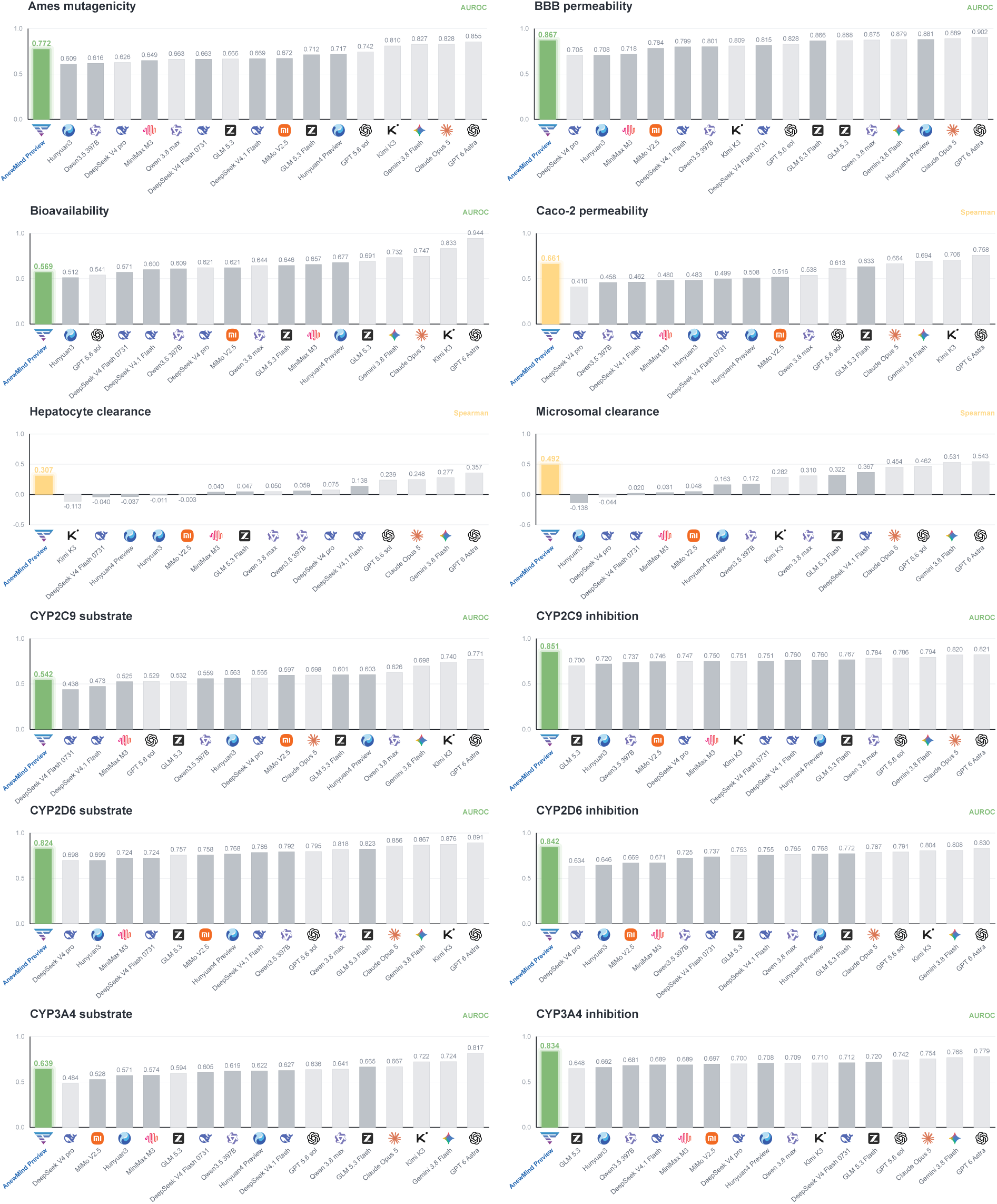
Performance of small molecule ADMET prediction II.

**Figure S4.**
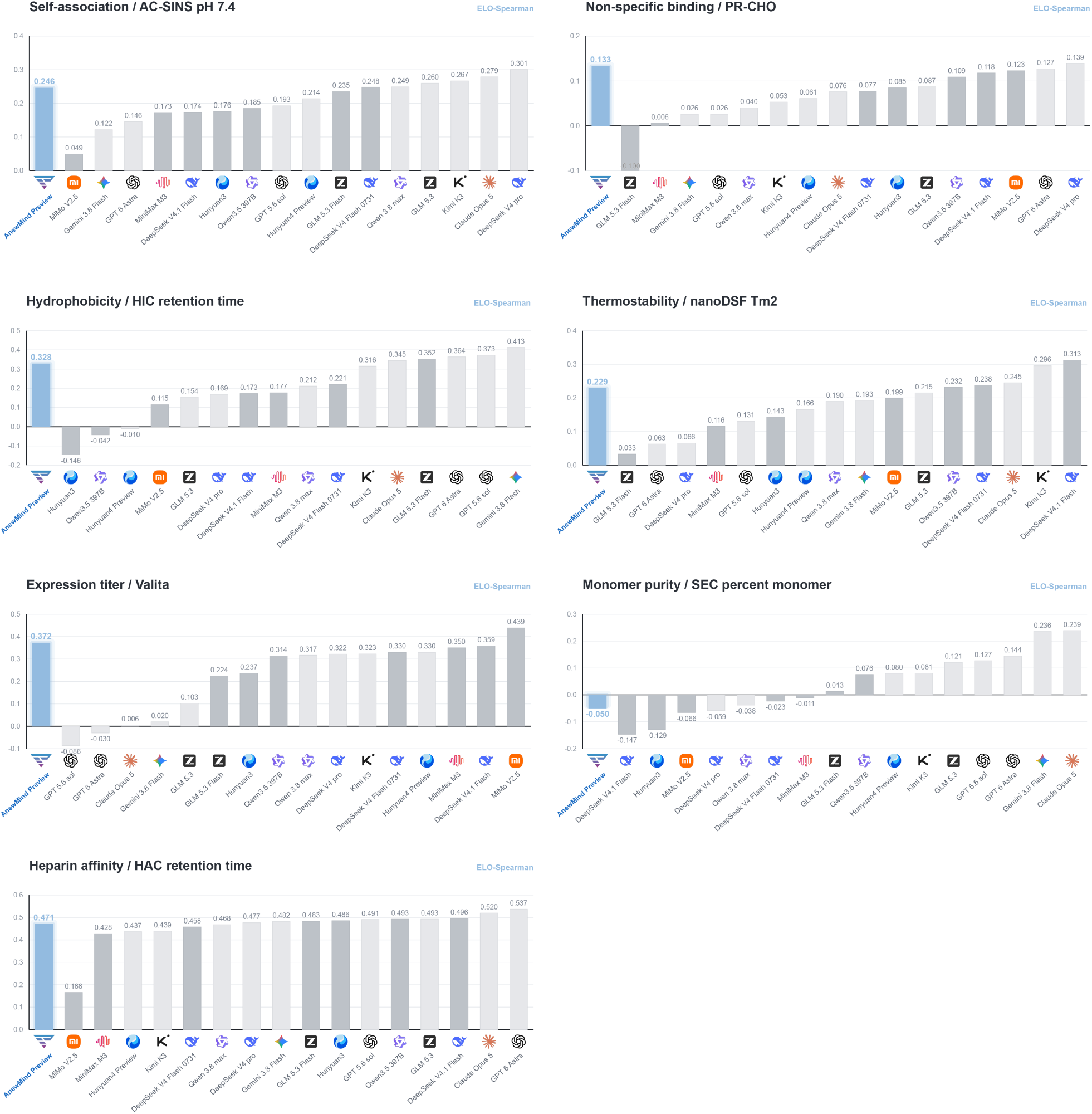
Performance of antibody developability prediction.

### A.7 ADMET Reasoning Case Studies from a Recent Patent

**Figure S5.**
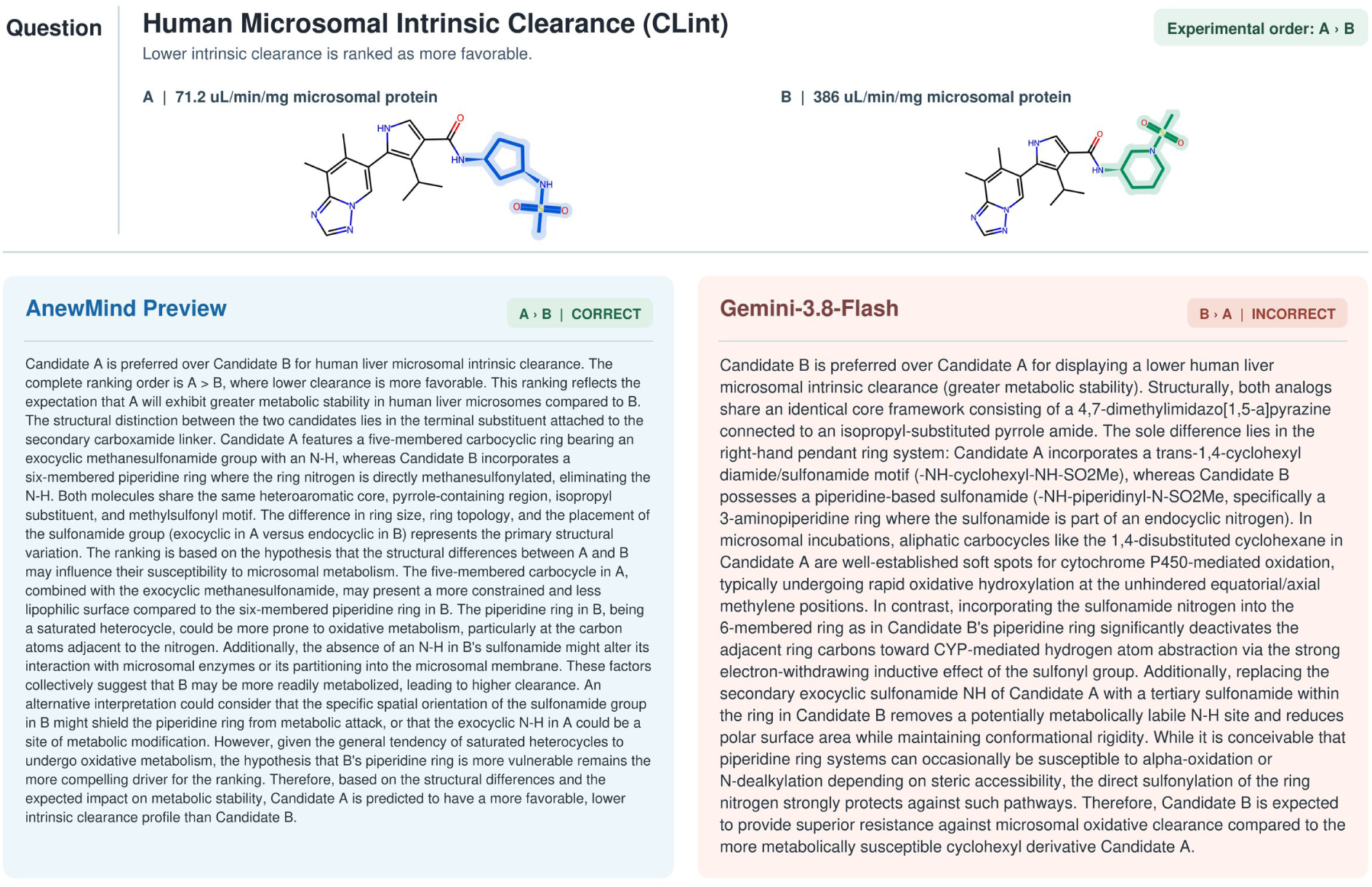
Human microsomal intrinsic clearance case study. AnewMind Preview correctly identified the experimental ranking and provided a structure-grounded ADMET rationale, whereas Gemini-3.8-Flash reversed the ranking.

**Figure S6.**
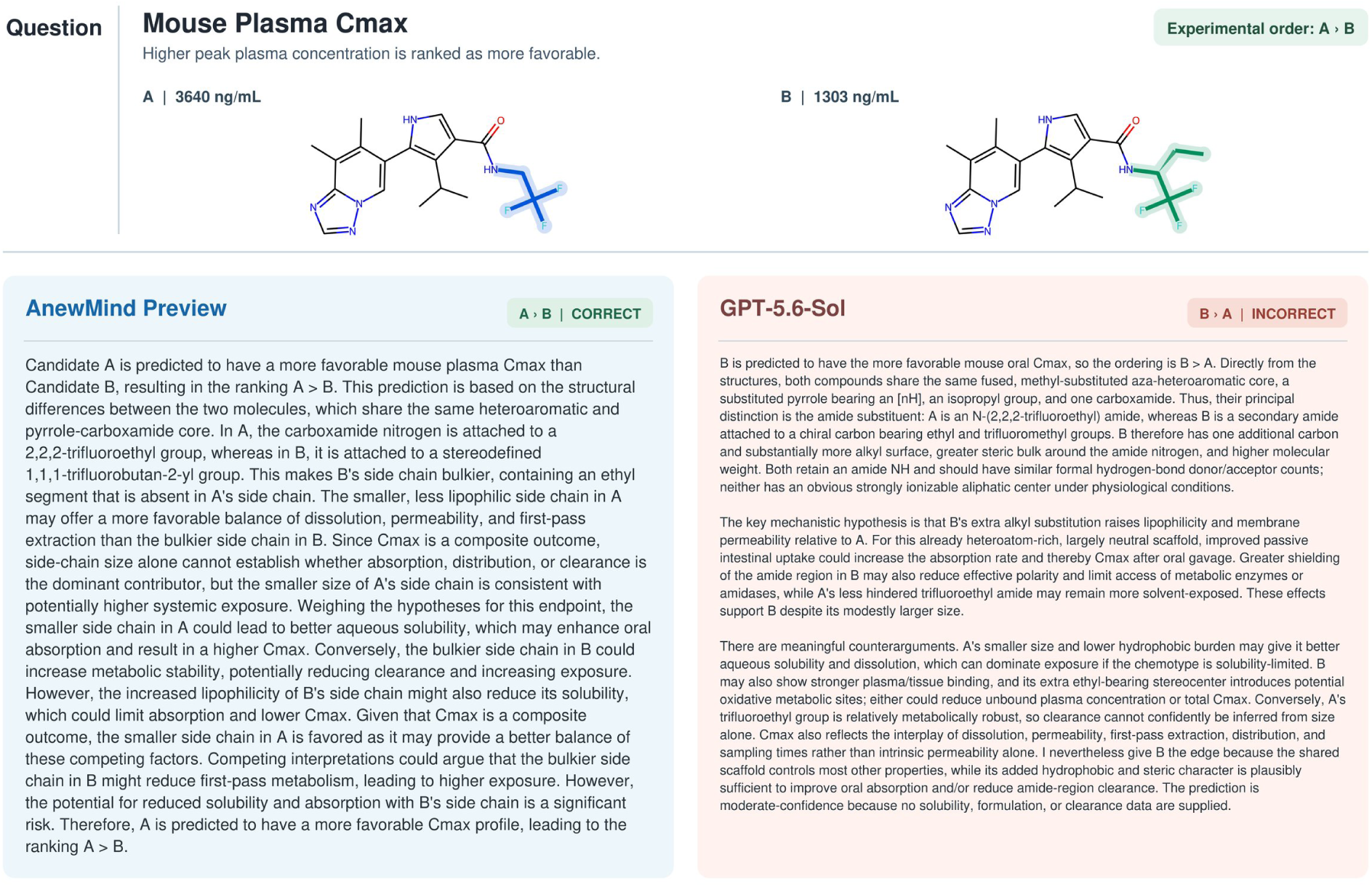
Mouse plasma Cmax case study. AnewMind Preview correctly identified the experimental ranking and provided a structure-grounded ADMET rationale, whereas GPT-5.6-Sol reversed the ranking.

**Figure S7.**
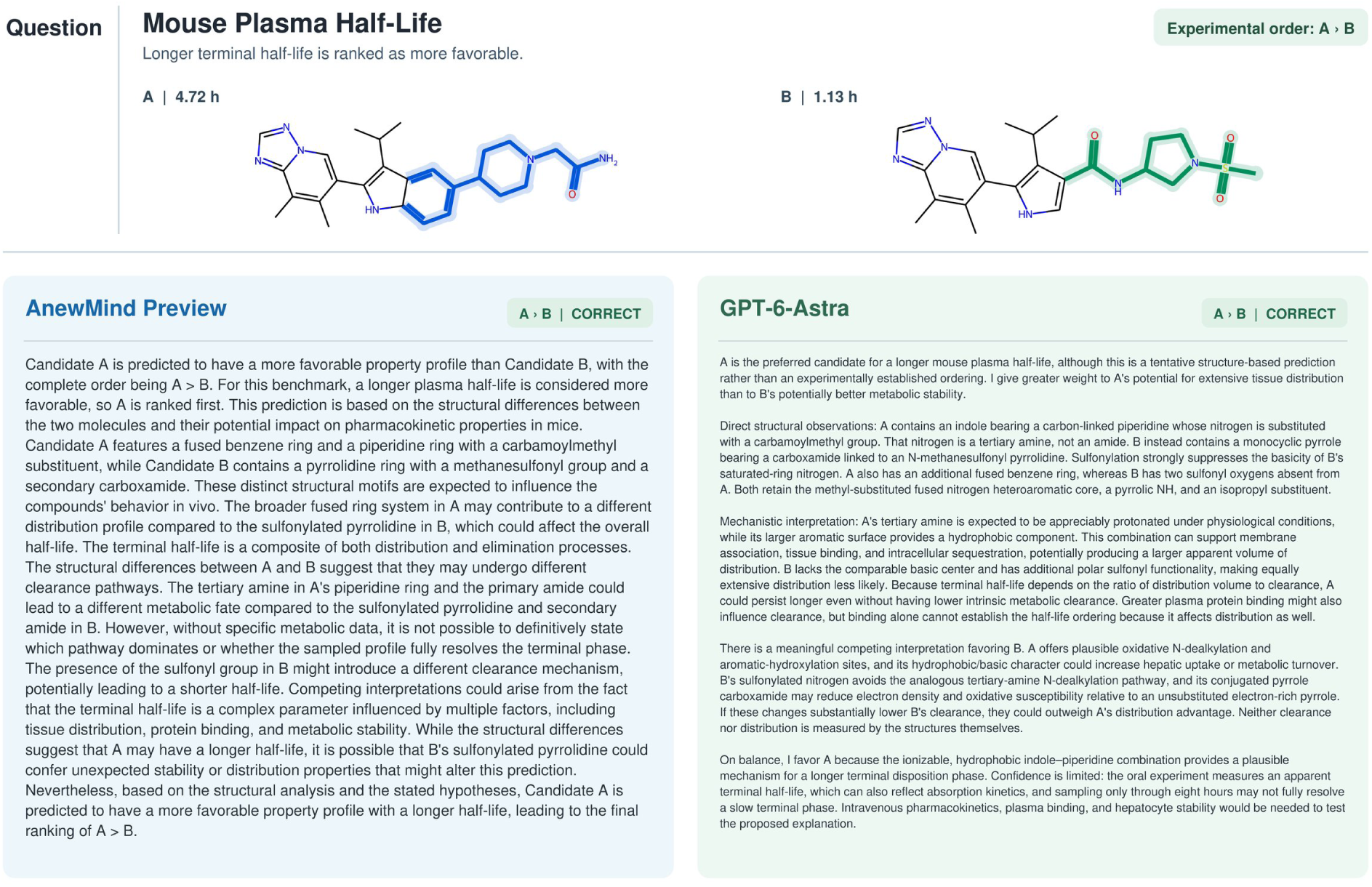
Mouse plasma half-life case study. Both AnewMind Preview and GPT-6-Astra correctly identified the experimental ranking and provided structure-grounded ADMET rationales.

### A.8 External-Model Evaluation Notes

#### Claude Opus 5 on PharmBench

Among the 200 PharmBench questions, Claude Opus 5 produced eight provider-side safety refusals. These responses contained no evaluable content and were therefore scored as incorrect.

#### Exclusion of selected GLM-5.3 results

GLM-5.3 is not shown in the small-molecule ADMET regression, GPQA Diamond, or MMLU-Pro comparisons because of task-specific output anomalies. Although isolated negative correlations were also observed for other models, GLM-5.3 produced negative correlations on six of the nine small-molecule regression endpoints, suggesting that it may not have consistently followed the requested ranking direction. In addition, 24 of 198 GPQA responses and 17 MMLU-Pro responses were truncated. The affected results were therefore excluded from the corresponding comparisons.

## Footnotes

1 The complete examples can be found here.

## Notes

### Competing Interest Statement

The authors have declared no competing interest.

